# Growth phase-specific gene regulation and algicidal interactions between a new *A. macleodii* strain and the model diatom *T. pseudonana*

**DOI:** 10.1101/2025.07.14.663959

**Authors:** David Wiener, Zinka Bartolek, Riley Dunklin, E. Virginia Armbrust

**Author notes:** corresponding author: David Wiener.

## Abstract

Phytoplankton-bacteria interactions are pivotal in marine ecosystems, influencing primary production and biogeochemical cycles. Diatoms, in particular, engage in diverse relationships with bacteria, ranging from mutualism to pathogenicity. However, the mechanisms governing the shift between these interactions and how they are shaped by host physiology and environmental context, remain unclear. To address this, we investigated how the diatom growth phase influences the interaction between a newly isolated *Alteromonas macleodii* strain from the Equatorial Pacific and the model diatom *Thalassiosira pseudonana*. We demonstrated that *A. macleodii*’s algicidal activity depends on the diatom’s growth phase, defensive capacity, and substrate availability. The algicidal effect manifests either during the diatom’s stationary phase or with an external source of organic carbon, implicating organic matter availability as a key driver. Transcriptomic analysis revealed that *A. macleodii* shifts from motility-associated to growth-associated gene expression patterns in response to the diatom’s growth phase and co-culture duration. Filtrate assays and fluorescence microscopy suggest a two-stage infection model: initial bacterial motility and exudate secretion induce diatom death, followed by bacterial aggregation around cellular debris. Comparative transcriptomics of *A. macleodii* with other algal hosts highlights host-specific bacterial responses, underscoring the context-dependent nature of these interactions. Together, these findings reveal how bacterial behavior and gene expression are modulated by host state and environmental cues, providing a molecular basis for the dynamic roles of diatom-bacteria interactions in shaping microbial community structure.

## Introduction

Phytoplankton exist in complex communities, sharing their microenvironment with diverse microbes, including heterotrophic bacteria [1]. Phytoplankton-bacteria interactions can influence algal physiology as well as large-scale processes such as carbon export and nutrient cycling [2]. Diatoms generate about 20% of global net primary production [3], and their ecological impact is shaped by interactions with marine bacteria that influence their growth, survival, and community composition [4]. Diatom-bacteria interactions range from pathogenic to mutualistic [4–6]. For example, certain bacteria provide diatoms with essential vitamins and growth-promoting compounds, while others release algicidal molecules that inhibit diatom growth or cause cell lysis. Transitions from mutualism to algicidal behavior within a given bacteria-algae pair have been demonstrated, driven by factors such as nutrient availability [7], co-culture duration [8–10], and community composition [11]. Diatoms can limit bacterial growth by secreting compounds like rosmarinic and azelaic acids, which promote beneficial bacteria while suppressing opportunists [12, 13]. Understanding the molecular mechanisms governing these interactions is crucial for elucidating the ecological role of diatoms, and the potential impacts of small-scale interactions on larger biogeochemical processes *Alteromonas macleodii* is a marine heterotrophic bacterium known for its versatility in forming both beneficial and detrimental interactions, highlighting its dynamic role in marine ecosystems [7, 14–17]. Notably, for a successful algicidal effect, some *A. macleodii* strains require physical attachment [7], while others rely solely on secreted compounds [15], demonstrating variable algicidal mechanisms. A recent study found that the *A. macleodii* behavior in co-culture with the model diatom *T. pseudonana* fluctuated from mutualistic to pathogenic with the addition of organic matter, which triggered bacterial movement toward the diatom and induced a protease-mediated algicidal effect. [7]. Yet, the molecular basis for the observed bacterial behavioral changes and how they adapt to exposure to diatoms in different physiological states remain hidden. Additionally, the *A. macleodii* strains tested in coculture with diatoms were obtained from coastal waters, so it remains unknown whether open-ocean strains exhibit similar interaction dynamics [7, 14, 16].

Growth phase shapes the physiological status of unicellular organisms and is often broadly divided into two main stages: the exponential phase, characterized by active growth, and the stationary phase, where cell division and death are balanced [18–20]. Diatoms show growth phase-specific transcriptomic [21], proteomic [22], and metabolomic [23] profiles. As diatom cells progress through their growth phases, they secrete organic matter, which serves as a substrate for bacteria [20]. These physiological changes of the diatom influence diatom-associated bacterial composition in laboratory settings and natural communities [23, 24]. Most co-culture experiments begin with diatoms in their exponential growth phase, leaving it unclear as to what extent encountering a diatom in a different growth phase influences the bacterial molecular response. Moreover, it remains uncertain how the outcome of specific species interactions depend on the diatom’s growth phase at the start of the co-culture.

Here, we characterize the interaction between a new *A. macleodii* strain recently isolated from the Equatorial Pacific Ocean and the model *T. pseudonana* through the diatom growth phases. We found that this *A. macleodii* strain exhibits an algicidal effect that depends on the diatom growth phase, organic matter availability, and a putative diatom defensive response. The new strain dynamically adjusted its transcriptional program over time in response to diatoms at different growth phases. Specifically, the *A. macleodii* strain downregulated genes related to motility (such as those involved in chemotaxis and flagellar assembly) and upregulated genes involved in cellular growth (including components of the translation machinery and ribosomal genes), depending on both the diatom’s growth phase and the duration of the co-culture. A combination of fluorescence microscopy and bacteria-free filtrate treatments suggests a two-stage infection model. In the first stage, bacteria swim and secrete harmful exudates that induce diatom death. This is followed by a second stage, where bacteria aggregate around diatom debris. Our study reveals that bacteria-algae interactions are context-dependent, with bacteria modulating their gene expression in response to host growth phases and environmental factors, shaping the interaction dynamics.

## Material & Methods

### Equatorial Pacific A. macleodii isolation & identification

The bacteria *A. macleodii* EP was isolated during the Gradients 4 cruise (TN397) in the Equatorial Pacific in December 2021 **(Supplemental Table 1)**. The bacteria was isolated from seawater collected at station 9 (140’W, 4.75’N) from a CTD cast at 15m depth. Approximately 1 mL of collected seawater was spread onto ½ YTSS plates (2g yeast extract, 1.5g tryptone, 20g sea salts, 15g agar in 1L of distilled water), and the plates were incubated for about 4 days at 20°C. Single colonies were picked and restreaked on ½ YTSS media plates until a pure culture was obtained. The identity of *A. macleodii* EP was determined by sequencing the 16S rRNA region (Sanger sequencing at Azenta) (**Supp. data 1**), and comparing the sequence to publicly available data via NCBI BLAST using the rRNA_typestrains/16S_ribosomal_RNA database. The 16S phylogenetic tree was generated using publicly available 16S sequences from NCBI that had high similarity to *A. macleodii* EP based on BLAST results **(Supplemental Table 2)**. Multiple alignment was done with MUSCLE [25], the aligned sequences were trimmed to be the same length in Geneious (Geneious Prime 2025.0.3), and the maximum likelihood tree was built using RaxML version 8.2.4 [26] (parameters:-m GTRGAMMA-f a-x 1-N 1000-p 1). The tree was visualized in Geneious.

### Co-culture experiments

The diatom *T. pseudonana* CCMP 1335 strain was maintained as a semicontinuous batch culture in 30 ml L1 + Si medium (NCMA) [27] at 20 °C with a 16:8 h light cycle, under ∼160 µmol photons m−2 s−1. In vivo chlorophyll a fluorescence was monitored (Turner Designs 10-AU fluorometer) over time as a proxy of cell division. *A. macleodii* EP glycerol stocks and other bacterias tested in this study **(Supplemental Table 1)** were streaked on marine agar plates (marine broth (MB) with 1.5% w/v agar) and grown at 25 °C for 48 hours. Single colonies were grown overnight in MB with shaking at 25 °C until an OD600 of ∼0.5 was reached (NanoDrop One Microvolume UV-Vis Spectrophotometer). The resulting bacterial cultures were diluted 1:10 and allowed to double twice (∼3:30 hours) before removing the media and washing twice with L1 + Si. The washed *A. macleodii* cells were added to a diatom culture at ∼40000 cells/ml final concentration. All experiments were performed in L1 + Si medium or L1 + Si medium supplemented with 2% MB (v/v) either at the beginning of the co-culture or at a specified time point.

### Exudates experiments

Samples were grown as previously described, and exudates were collected by filtering twice through a 0.22 µm glass fiber filter at the specified time points. Bacteria were cultured as described before being inoculated into the exudates at a concentration of 40000 cells/ml. Diatoms from exponentially growing cultures were diluted into the various exudates at 2% (v/v), initiating the experiments with an initial concentration of approximately 200,000 cells/ml.

### Flow cytometry

Samples were incubated with SYBR Green (1:5000) for 30 minutes at room temperature in the dark, followed by analysis using a Guava easyCyte 11HT Benchtop Flow Cytometer. *T. pseudonana* cells were detected based on chlorophyll autofluorescence (emission at 642 nm) and forward scatter, and *A. macleodii* EP cells were detected by SYBR fluorescence (emission at 532 nm) and forward scatter. Gating parameters were established using control cultures for each species.

### Nutrients (PO4, Si(OH)4, NO3, NO2, NH4) measurements

Samples were filtered through a 0.22 µm glass fiber filter, immediately frozen, and were analyzed at the UW Marine Chemistry Laboratory with a Seal Analytical AA3, following the protocols of the WOCE Hydrographic Program [28].

### Dissolved Organic Carbon (DOC) measurements

Samples were filtered using 25 mm carbon-cleaned GF/F filters (combusted at 450°C for 4 hours), immediately frozen, and were analyzed at, the UW Marine Chemistry Laboratory with a Shimadzu TOC-Vcsh DOC analyzer, following the protocols of the WOCE Hydrographic Program [28].

### RNA extraction and library preparation

Total RNA was extracted from 0.2 µm polycarbonate membrane filters using the Zymo Direct-zol RNA MiniPrep Plus kit for all samples. For T. pseudonana transcriptome ribodepletion calibration, three samples obtained from exponentially growing cells were prepared: Total RNA, poly(A) selected with oligo dT-beads (Dynabeads mRNA DIRECT Kit, Life Technologies), and ribodepleted using a tailored Thalassiosira pseudonana riboPOOL following the manufacturer’s instructions. For Duo-RNA-seq, samples were prepared in triplicate for each time point and ribodepleted using the *T. pseudonana* riboPOOL combined with the Pan-Bacteria riboPOOL, with beads mixed in a 1:1 ratio. At this point, libraries were submitted for preparation and sequencing to the Northwest Genomics Center (University of Washington) using a NextSeq (Illumina) platform.

### Ribo-deplition efficiency calculation

Sequence reads were trimmed using Trimmomatic 0.39 [29], run in paired-end mode with the adaptor and other Illumina-specific sequences (ILLUMINACLIP) set to TruSeq3-PE.fa:2:30:10:1, leading and trailing quality thresholds of 25, a sliding window trimming approach (SLIDINGWINDOW) of 4:15, an average quality level (AVGQUAL) of 20, and a minimum length (MINLEN) of 60. For RNA identification, paired-end reads were aligned with STAR v2.7.10b [30] to the *T. pseudonana* genome (Thaps3, FilteredModels2) [31] plus an additional “chromosome” containing an updated version of the ribosomal sequences determined using Nanopore sequencing [32]. Default parameters were used, with the maximum intron length limited to 500 nucleotides (‘–alignIntronMax 500’). Reads that aligned to both the *T. pseudonana* genome and the rRNA chromosome were assigned to the latter. Reads per chromosome were calculated using the samtools idxstats command [33] and assigned to gene models using the featureCounts tool [34]. The rRNA read fraction was calculated as the total number of reads assigned to the rRNA chromosome divided by the total aligned reads. Gene models read fraction was calculated as the total number of reads assigned to a gene model divided by the total number of aligned reads.

### A. macleodii genome determination

To determine the most similar genome option, we first selected all *A. macleodii* genomes available in the NCBI genome database that met the following criteria: A) Complete assembly level, B) Annotated, C) Not from metagenome-assembled genomes (MAGs), and D) Not labeled as atypical. This resulted in a total of 14 genomes **(Supplemental Table 3)**. Paired-end reads were analyzed as previously described but mapped to the 14 genomes in parallel. To determine which genome best matched the *A. macleodii* EP strain, we used three metrics: the percentage of uniquely aligned reads, the fraction of reads assigned to gene models, and the number of gene models with more than 5 reads.

### Duo-RNA-seq analysis

Paired-end reads were analyzed as previously described, but mapped to a combined T. pseudonana (Thaps3, FilteredModels2) and A. macleodii Te101 genome [31, 35]. Gene counts for each genome were processed separately, with trimmed mean of M-values (TMM) normalization and RPKM calculation performed using the EdgeR package. [36]. The batch effect from different rounds of ribodepletion was removed using the ComBat function [37]. For PCA analysis, counts were scaled and then analyzed using the R function prcomp. Differential expression analysis was conducted using the limma package [38] for the specific contrasts. To identify significant Gene Ontology (GO) terms associated with differentially expressed genes, we employed the topGO R package [39] using GO annotations provided by Hou et al [35]. Additionally, KEGG Orthology (KO) IDs for each gene model were identified using the BlastKOALA tool [40], and selected pathways were retrieved manually. Additionally, enrichment of KOG categories among *T. pseudonana* differentially expressed genes was tested using a proportion test (Chi-squared test for equality of proportions) against their distribution in the full transcriptome.

### Fluorescent microscopy

200 µl of *T. pseudonana* cultures or co-cultures with *A. macleodii* EP were stained with SYBR Green (1:5000) and incubated for 30 minutes at room temperature in the dark. After incubation, samples were centrifuged at 10,000 rpm for 1 minute, and the supernatant was discarded, leaving a final volume of 10 µl. The concentrated samples were then mounted onto transparent slides, and images were captured at 20x magnification using a Leica DMi8 microscope. Fluorescent images were taken for DNA (green channel), chlorophyll (red channel), and brightfield. Image processing was performed using ImageJ [41].

### Casein Plate assay

Casein agar plates were prepared in small batches. For every 100 mL of artificial seawater, 0.3 g of casein protein was well dissolved. After full dissolution, 0.2 g of yeast extract and 1.5 g of Bacto-agar were added. After autoclaving, plates were stored in the fridge until ready for use. Two biological replicates and two technical replicates were prepared for the following conditions: diatom exudate (*T. pseudonana* grown 24h in L1+Si medium), co-culture exudate (*T. pseudonana* and *A. macleodii* grown 24h in L1 + Si medium), bacteria exudate (*A. macleodii* grown 24h in L1+Si medium supplemented with 2% MB (v/v)), and co-culture+MB exudate (*T. pseudonana* and *A. macleodii* grown 24h in L1 + Si medium supplemented with 2% MB (v/v)). Growth conditions and filtration were performed as previously described. Immediately after filtration, 7 μL droplets of each condition were placed onto marked sections of the casein agar plates. Plates were stored in the same conditions as the co-culture experiments. The diameter of each halo was measured six days after plating.

### Statistical Analysis

To evaluate the effects of experimental factors on the cell numbers of either T. pseudonana or A. macleodii, while accounting for variability across replicates, we applied a generalized additive mixed model (GAMM) approach using the gamm function from the mgcv package in R [42]. Fixed effects included the relevant experimental conditions and their interactions, while replicate identity was treated as a random effect to account for the repeated measures. An analysis of variance (ANOVA) was performed on the GAMM model to assess the significance of the fixed effects. Post hoc pairwise comparisons were conducted using estimated marginal means (EMMs) with Bonferroni-adjusted p-values to control for multiple testing, as implemented via the emmeans package [43]. Additionally, the Wilcoxon signed-rank test was applied to compare treatments, where appropriate.

### Data availability

The data sets generated in this study for RNA expression are available on GEO under the accession number GSE291849 *(reviewer token: gxyniuouvfydfkh)*. *A. macleodii* MIT1002 co-culture with *Prochlorococcus* RNA expression data was obtained from GEO (accession number GSE73511).

## Results

### Characterization of the interaction between A. macleodii EP and T. pseudonana across different growth phases

We cultured a new *A. macleodii* isolate from seawater collected on a research cruise (Gradients 4) in the Equatorial Pacific (0.5 N, 139.73 W, 15 m Depth on December 5, 2021). The new isolate displayed 16S ribosomal DNA sequence identity with publicly available 16S rDNA sequence for two *A. macleodii* strains **(Fig. S1, Supplemental Table 2)**. We designate this new isolate as *A. macleodii* EP. Given the known variety of interactions of *A. macleodii* with unicellular algae, we co-cultured the new *A. macleodii* isolate with the model diatom *T. pseudonana* at early exponential, mid-exponential, and stationary phases, without the addition of an external organic carbon source to the media. We defined the *T. pseudonana* stationary phase as 7 days after the last cell doubling when the cell concentration reached ∼800,000 cells/ml. The two exponential growth phases were defined relative to stationary phase: Early exponential phase cultures were at ∼25,000 cells/ml, approximately five doubling events before reaching stationary phase cell concentrations and mid-exponential/exponential phase cultures were at ∼200,000 cells/ml, two doubling events before reaching the stationary phase cell concentration **(Fig. 1A-B, lower panels)**. When co-cultured with diatoms in early-or mid-exponential phase, *A. macleodii EP* abundance increased over the first few days and then decreased in abundance, likely due to depletion of available organic carbon **(Fig. 1A upper panel)**. In contrast, when *A. macleodii* EP was co-cultured with a stationary phase *T. pseudonana* culture, *A. macleodii EP* reached significantly higher cell concentrations (ANOVA followed by post hoc pairwise comparisons with Bonferroni-adjusted p-values: Early Exponential - Mid Exponential=1, Early Exponential - Stationary 0.0001, Mid Exponential - Stationary <.0001), presumably due to increased availability of organic carbon released by *T. pseudonana* over the growth cycle **(Fig. 1A, upper)**. During the stationary phase co-culture, *A. macleodii* displayed an algicidal effect on *T. pseudonana,* resulting in a 0.3-fold decrease in final diatom abundance **(Fig. 1A, lower panel)**. These results suggested that either stationary phase *T. pseudonana* cells were particularly vulnerable to *A. macleodii EP* or that a sustained increase in *A. macleodii EP* abundance was required for the algicidal impact.

**Figure 1.**
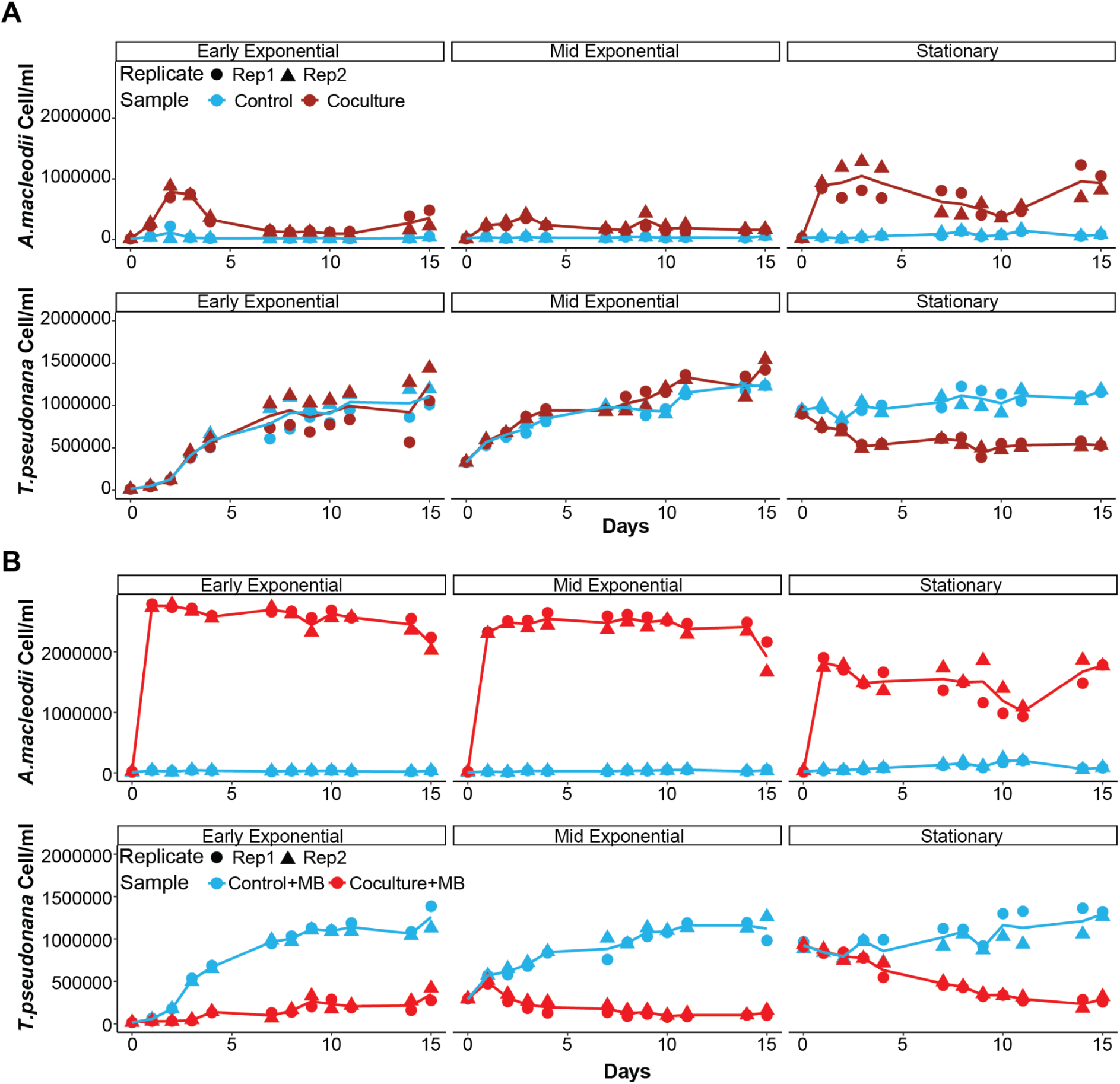
Initial culture conditions for *T. pseudonana* play a key role in determining interaction dynamics with *A. macleodii* EP. **(A)** Growth curves of bacterial cells (top) and diatom cells (bottom) when grown in co-culture (magenta symbols) or as mono-cultures (blue symbols); replicates indicated with different symbol types (circles, triangles), with the line indicating the average of two replicates. Each facet corresponds to a different diatom growth phase at the start (t=0 days) of the experiment (early exponential, mid-exponential, or stationary phase). **(B)** Similar to (A), but with the addition of 2% (v/v) marine broth at the start of the experiment (red symbols).

To evaluate whether stationary-phase *T. pseudonana* cells displayed enhanced sensitivity to *A. malceodii* EP, we controlled bacterial abundance in each set of co-cultures by the addition of 2% (v/v) marine broth (MB) as an external organic nutrient source. With the addition of organic carbon, *A. macleodii* EP reached high cell concentrations in each co-culture. Regardless of the diatom growth phase, the supplemented co-cultures with *A. macleodii* EP either prevented an increase in *T. pseudonana* abundance (early exponential) or resulted in a decrease in *T. pseudonana* abundance (mid-exponential and stationary phase) over time **(Fig. 1B)**. A similar algicidal effect was observed if the organic carbon was added 4 days after initiation of the co-culture **(Fig S2A)**. Two other bacteria - *Ruegaria pomeroyii* and *Sulfitobacter sp* did not display an algicidal impact under the same co-culture conditions, confirming that the observed *A. macleodii* EP algicidal effect was not an artifact of our culture conditions **(Fig. S2B)**. Unexpectedly, the final concentration of *A. macleodii* EP in the amended co-cultures varied depending on the initial growth phase of *T. pseudonana*. When co-cultured with stationary-phase *T. pseudonana*, *A. macleodii* EP cell numbers were significantly reduced by 0.66-fold by the 24-hour time point **(Fig. S2C)** compared to those co-cultured with exponential-phase *T. pseudonana*. Similarly, a 0.79-fold reduction was observed when MB was added four days after the initiation of late-exponential-phase *T. pseudonana* co-cultures **(Fig. S2D)**. These results suggest that *T. pseudonana* somehow suppressed *A. macleodii* EP growth in a manner dependent on diatom cell concentration.

### Diatom’s defensive response limits A. macleodii EP growth

To evaluate whether *T. pseudonana* differentially released compounds over the growth cycle that either supported or suppressed *A. macleodii* EP growth, we grew axenic mono-cultures of *T. pseudonana* and collected cell-free culture media (exudate) from mid-exponential and stationary phases. Final *A. macleodii* EP cell abundances were 2.4-fold higher when grown in the stationary phase exudate than in the mid-exponential exudate, consistent with the assumption that organic matter released by *T. pseudonana* into the growth media supported the growth of *A. macleodii EP* **(Fig. 2A)**. We compared these final cell abundances with those obtained in co-cultures with or without the MB supplement of organic matter. During the mid-exponential phase of *T. pseudonana*, final *A. macleodii* EP concentrations were lowest in the unsupplemented co-culture and highest in the MB-supplemented co-culture, with the *T. pseudonana* mono-culture exudate resulting in an intermediate concentration of *A. macleodii* EP **(Fig. 2A, left panel)**. In contrast, the stationary phase mono-culture exudate supported the highest final concentration of *A. macleodii*, with lower final concentrations in both the supplemented and unsupplemented co-cultures **(Fig. 2A, right panel)**. These results suggest that in co-culture, growth of *A. macleodii* EP was limited by factors beyond availability of organic matter, and that attributes of *T. pseudonana* physiology across the different growth phases actively limited growth of *A. macleodii* EP.

**Figure 2.**
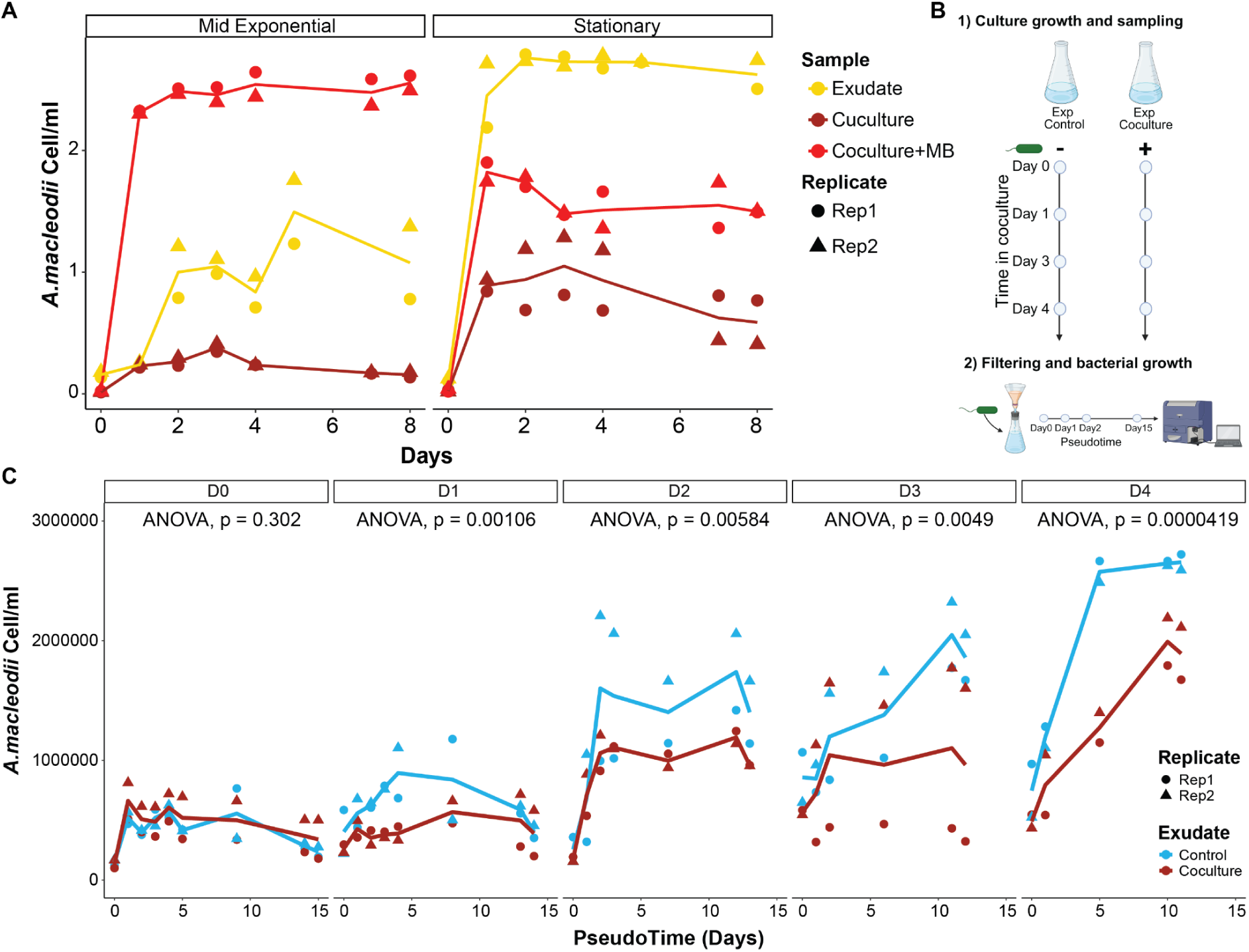
*T. pseudonana* imposes constraints on bacterial growth beyond nutrient limitation. **(A)** Scatter plot of bacterial cell counts measured by flow cytometry, with lines indicating the mean of two replicates. Colors distinguish bacterial growth in co-culture (magenta), in co-culture with Marine Broth (co-culture+MB) (red), and with exudates from diatom cultures (yellow). Each facet corresponds to a specific growth phase. **(B)** Schematic representation of control and co-culture filtrate experiments. **(C)** Scatter plot showing bacterial cell counts in each filtrate. Colors distinguish filtrates obtained from control (blue) or co-culture (magenta) conditions. ANOVA p-values from mixed effects models were calculated to quantify the effects of the exudate treatment. Facets indicate the day of culture when the filtrate was collected.

To further investigate the apparent defensive response of *T. pseudonana* and distinguish between a constitutive effect and one triggered by the presence of bacteria, we compared bacterial growth in exudates generated from either *T. pseudonana* mono-cultures or co-cultures of *T. pseudonana* and *A. macleodii* EP collected at different times after initiation of the mono-cultures or co-cultures **(Fig. 2B-Fig S3A)**. We reasoned that, provided sufficient organic matter was present to support bacterial growth, *A. macleodii* EP would reach higher abundances in exudates from *T. pseudonana* mono-cultures than in exudates from co-cultures if, indeed, the presence of bacteria triggered a defensive response in *T. pseudonana*. Immediately after the start of the co-cultures, there were no significant differences in *A. macleodii* EP abundances **(Fig. 2C, facet D0)**. However, after 24 hours, the mono-culture exudates supported higher bacterial abundances than the co-culture-derived exudates **(Fig. 2C, facet D1)**. Moreover, the differences between growth in mono-culture and in co-culture exudates increased in subsequent time points (**Fig 2C, facets D2, D3, D4**). The rate of draw-down of inorganic nutrients was comparable between the mono-cultures and co-cultures, indicating that *T. pseudonana* grew similarly under both conditions **(Fig S3B)**. Accumulation of DOC was greater in the mono-culture than in the co-culture, presumably because *A. macleodii* EP drew down DOC for growth. The continued accumulation of DOC in the co-culture, however, suggested that *A. macleodii* EP was unable to take up the DOC at the same rate as the rate of DOC production by *T. pseudonana*. Furthermore, the maximal cell abundance attained by *A. macleodii* EP was significantly higher when the bacteria were grown on the mono-culture exudate than on the co-culture exudate, suggesting that the bacteria were unable to utilize available DOC and attained lower cell numbers on the co-culture exudate **(Fig. 2C-Fig. S3C)**. There was no significant correlation between the ratio of maximum bacterial abundance in mono-cultures and co-cultures and the ratio of organic matter in the two conditions (R=0.3, p=0.47) **(Fig. S3D)**. Thus, the defensive response is triggered by the presence of bacteria and remains in the liquid phase of the culture after cell filtration, suggesting either the secretion of a defensive compound or changes in culture conditions as potential mechanisms.

Together, these observations suggested that this interaction was governed by a bacterial algicidal effect, the diatom growth phase, availability of organic matter, and a diatom-triggered defensive response. During the *T. pseudonana* stationary phase, organic matter secretion enabled *A. macleodii EP* to reach cell numbers sufficient to cause diatom death. In contrast, during the exponential phase, *T. pseudonana* appeared to limit bacterial growth through the secretion of defensive compounds, despite adequate availability of organic matter. Addition of marine broth bypassed the diatom defensive effect, allowing bacterial proliferation and algicidal effects.

### Duo RNA-seq reveals the gene expression regulatory landscape of the A. macleodii EP–T. pseudonana interaction

To investigate regulatory dynamics of gene expression in *A. macleodii* EP and *T. pseudonana* during their interaction across growth phases, we measured transcriptional profiles at 30 minutes, 36 hours, and 8 days of co-culture **(Fig. 3A)**. Use of a specific *T. pseudonana* riboPOOL kit designed based on an updated version of the rRNA sequences [32] minimized host rRNA representation in the sequence data **(Fig. S4A-B)**. We aligned the resulting sequence reads to the *T. pseudonana* genome and to 14 *A. macleodii* genomes. The greatest number of reads aligned to the *T. pseudonana* and the *A. macleodii* Te101 genomes [35] **(Fig. S4C).** Moreover, we confirmed the ability to distinguish *A. macleodii* and *T. pseudonana* reads by analyzing the control cultures, which showed no bacterial reads, and by detecting an increase in bacterial reads over time, particularly in the stationary-phase co-culture **(Fig. S4D)**.

**Figure 3.**
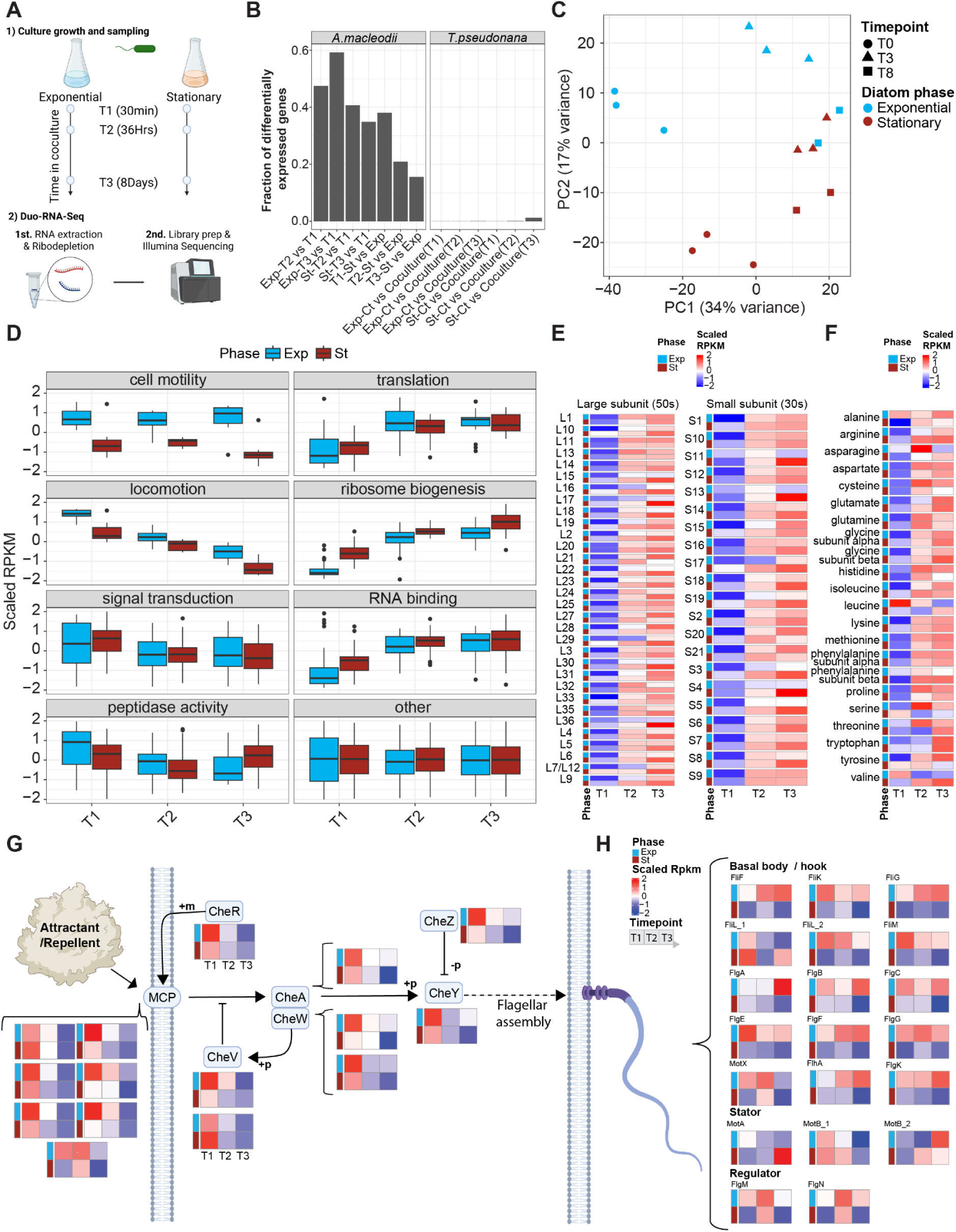
Gene expression regulatory landscape in the *A. macleodii* EP–*T. pseudonana* interaction. **(A)** Schematic representation of the sampling design and Duo-RNAseq library preparation. **(B)** Bar plot showing the fraction of differentially expressed genes in both genomes for specific comparisons. **(C)** Principal component analysis (PCA) using scaled RPKM reads from *A. macleodii* EP. **(D)** Boxplot showing scaled RPKM values for selected GO groups in A. macleodii EP across time points and host growth phases: exponential (blue) and stationary (magenta). RPKM values are averaged across replicates. **(E)** Heatmap showing scaled RPKM values for ribosomal proteins identified using the KEGG BLAST KOALA tool and assigned to the Ribosome KEGG pathway (ko031011). **(F)** Similar to (E), but for aminoacyl-tRNA synthetases from the Aminoacyl-tRNA biosynthesis KEGG pathway (ko00970). **(G)** Schematic representation of gene expression changes in the Bacterial Chemotaxis (ko02030) KEGG pathway. Heatmaps were generated from the scaled average RPKM values between replicates. Row annotations distinguish between exponential (blue) and stationary (magenta) samples, while each column represents a time point (T1, T2, T3), organized from left to right. Gene names are indicated above each heatmap. The dashed arrow illustrates how chemotaxis signaling, mediated by methyl-accepting chemotaxis proteins (MCPs), controls flagellar assembly in response to environmental signals. The “attractant/repellent blob” represents the chemical gradients sensed by MCPs, which modulate bacterial movement. MCPs are located in the cell membrane, where they interact with chemical signals, while the flagella, also in the membrane, are responsible for bacterial swimming. **(H)** As in (G), but for the Flagellar Assembly (ko02040) KEGG pathway. Genes were divided into the categories Basal body/hook, Stator, and Regulator.

To explore the differences triggered by bacterial presence and the effects of the diatom growth phase and time in co-culture, we identified differentially expressed genes in the corresponding pairwise comparisons **(Fig. 3B-Fig. S5A)**. Interestingly, the presence of the bacteria did not elicit a significant response in the diatom transcriptome, with only ∼90 (0.7%) differentially expressed *T. pseudonana* genes **(Fig. 3B-Fig. S5A)**. PCA analysis revealed that transcript abundance profiles of exponential-phase samples progressed through the growth phases independently of bacterial presence **(Fig. S5B)**, suggesting that the presence of bacteria did not significantly alter the growth profile of the diatom at the transcriptional level.

In contrast, during co-culture, 20 to 40% of *A. macleodii* EP genes (200-600 genes) were differentially expressed, depending on the sampling time point **(Fig. 3B-Fig. S5A)**. PCA-based analyses indicated that ∼34% of the variability was described by the first principle component (PC1) that followed the progression of time in co-culture; ∼16% of the variability was described by PC2 that separated bacteria co-cultured with *T. pseudonana* in exponential versus stationary phase **(Fig. 3C)**. The most significant differences occurred at the earliest time point (T1), suggesting that bacteria rapidly sensed and responded to their environment. Overall, these results suggest that the progression of the diatom culture and the initial conditions of the co-culture shape the outcome of the interaction and the transcriptome of *A. macleodii* EP.

A gene ontology (GO) enrichment analysis of differentially expressed *A. macleodii* EP genes revealed consistent changes across time when co-cultured with either stationary-phase or exponential-phase diatoms **(Fig. S5C)**. At early time points, upregulated genes were enriched for Biological Process GO terms related to cell motility, locomotion, and signal transduction, as well as Molecular Function GO terms associated with peptidase activity. At later time points, enriched Biological Process GO terms included translation and ribosome biogenesis, while Molecular Function terms were related to RNA binding. Similar GO terms were differentially expressed when comparing bacteria exposed to stationary-versus exponential-phase diatoms **(Fig. S5C)**. The genes categorized with the GO terms locomotion and signal transduction were downregulated over time in bacteria co-cultured either with exponential-phase or stationary-phase diatoms **(Fig. 3D)**. Conversely, the genes categorized with the GO terms ribosome biogenesis, translation, and RNA binding were upregulated over time in bacteria co-cultured either with exponential-phase or stationary-phase *T. pseudonana* **(Fig. 3D)**. Moreover, fold change differences across GO groups were significantly higher at T1—except for cell motility—suggesting a rapid transcriptional adjustment to environmental conditions **(Fig. S5D)** Notably, genes related to cell motility were consistently downregulated when co-cultured with stationary-phase *T. pseudonana*, indicating a prolonged effect of the initial co-culture environment **(Fig. 3D)**. These dynamics were not observed in the distribution of other GO groups, highlighting the specificity of the response **(Fig. 3D)**.These transcriptional profiles appear to be quantitative, with transcript abundances transitioning progressively over time from low to high (or vice versa), suggesting a graded adaptation to the diatom-derived environment **(Fig. 3D)**. This reflects a gradual shift from a motile sensing profile, where *A. macleodii* secretes peptidases as part of its response to environmental cues, to a growth-oriented profile.

The progression of gene expression between exponential and stationary co-cultures differed for genes with predicted peptidase activity. In exponential co-cultures, transcripts levels for the peptidase-encoding genes decreased consistently across all time points, whereas in stationary co-cultures, transcript levels initially decreased but recovered to T1 levels by T3. Additionally, at the first two time points, expression was significantly higher in bacterial co-culture with exponential-phase diatoms, while at T3 the relationship was reversed **(Fig. 3D)**. Furthermore, peptidase predicted genes transcripts levels changed not only in magnitude over time, but also in which specific peptidases were expressed **(Fig. S6A, left)**. Similar changes across time and stages in transcript levels were detectable for genes encoding *A. macleodii* EP proteases **(Fig. S6A, right)**. Together, these results demonstrate that both the quantity and identity of transcripts associated with peptidases and proteases are influenced by time in co-culture and the physiological state of the diatoms.

Three general categories of differentially transcribed genes illustrate the transition between different bacterial functions over time and in response to the environment. One category includes genes encoding ribosomal proteins from both the large and small subunits, as well as tRNA synthetases. Transcript abundances for these genes increased over time in all co-co-cultures and were lower overall in bacteria facing exponential diatoms, especially at the time point T1 **(Fig. 3E)**. Similarly, transcripts for 18 of the 22 detectable aminoacyl-tRNA synthetases were upregulated with time in co-culture and 15 of the 22 were in higher concentrations when co-cultured with stationary diatoms at T1 **(Fig. 3F)**. Ten of the 22 remained at higher abundances when co-cultured with stationary diatoms at T3. The second category of differentially expressed genes included those involved in chemotaxis signaling. These genes were downregulated through time in co-culture regardless of whether the diatoms were in exponential or stationary phase **(Fig. 3G)**. This subset of genes include those associated with key signaling components such as the methyl-accepting chemotaxis proteins (MCPs), which reside in the bacterial membrane and act as primary sensors of environmental chemical gradients. Also included are genes that encode signal transduction proteins, such as the CheA kinase, motor regulators like CheY and CheZ, adaptation-related proteins such as CheR, and coupling proteins like CheV and CheW. The third category of genes encode flagellar proteins. Most flagellar genes diverged between exponential and stationary phase conditions **(Fig. 3H)**. Genes encoding the basal body and hook structures were significantly higher when co-culture the exponential phase diatom and lower when co-culture with the stationary phase diatom. Genes encoding the flagellar stator complex (such as MoA and MA) and flagellar regulator (such FlgM and FlgN) proteins displayed gene-specific transcriptional patterns. Specifically, one paralog of MoB was downregulated over time, while the other was upregulated, suggesting differential regulation. The expression of MobA remained stable throughout the experiment. The expression patterns of FlgM and FlgN did not show clear differences between stationary and exponential diatom growth phases, indicating their regulation is less affected by these growth conditions. In summary, despite changes in chemotaxis signaling mRNA levels, *A. macleodii* EP maintains upregulation of flagellar genes when exposed to exponential-phase *T. pseudonana* cells, but shows reduced expression in response to stationary-phase cells. These findings suggest that *A. macleodii* EP adjusts the balance between motility and growth through transcriptomic changes during co-culture, depending on the diatom’s growth phase. This pattern aligns with a behavioral shift from searching for to exploiting a nutrient source. The transition is accompanied by dynamic changes in peptidase and protease expression—both in magnitude and specificity **(Fig. 3D, Fig. S6A)**—indicating that *A. macleodii* modulates its motility and enzymatic machinery in response to the physiological state of *T. pseudonana* to optimize nutrient acquisition.

### A. macleodii EP algicidal effect is mediated via the secretion of a algicidal compound

The transcriptional data suggested that *A. macleodii* EP transitions from chemotactic swimming behavior during the early phase of co-culture when bacterial abundance is low to algicidal impacts coincident with production of distinct proteases and peptidases. To determine whether *a* direct cell-cell contact was required for the algical effect, we examined the phenotypic behavior of *A. macleodii* EP in the different co-cultures over time. We hypothesized that if close-range interactions were necessary for algicidal activity, then *A. macleodii* EP should cluster around *T. pseudonana* cells during the later stages of co-culture. Co-cultures were prepared using diatoms in exponential and stationary phases, with or without MB supplementation. After 3 days in co-culture at peak algicidal activity, *A. macleodii* EP cells were detectable by microscopy **(Fig. 1A–Fig. 4A)** in both the stationary phase co-cultures and in those supplemented with MB. At this early time point, the bacteria appeared randomly distributed, with no clear association with *T. pseudonana* cells **(Fig. 4A-left)**. By day 8 when diatom cell number was significantly reduced, the distribution pattern shifted: *A. macleodii* EP cells clustered around apparent diatom debris **(Fig. 4A-right)**. This spatial reorganization suggested that the bacteria were actively interacting with the lysed diatom material, potentially breaking it down for necessary substrates.

**Figure 4.**
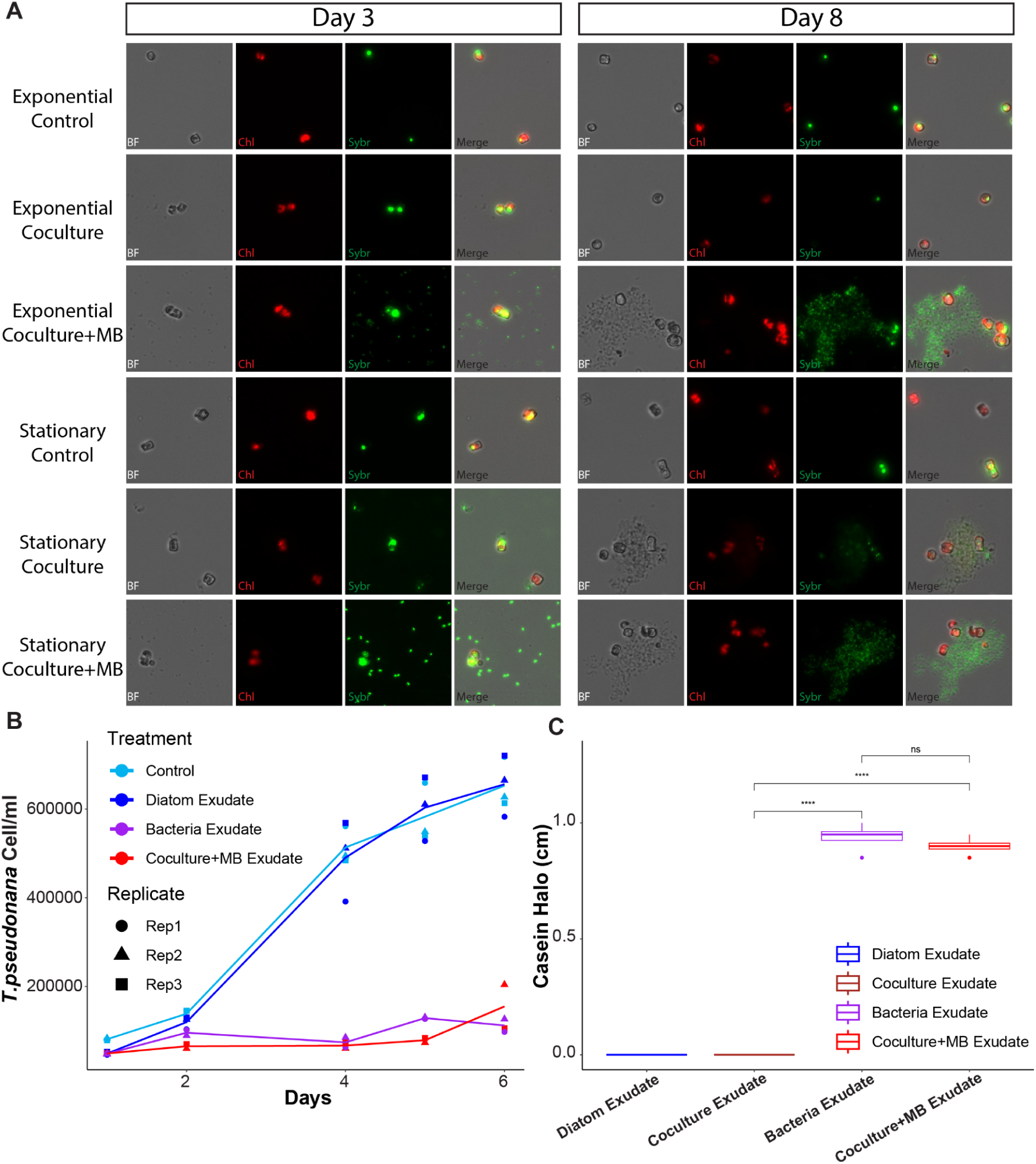
Bacterial infection occurs in two stages, likely mediated by the secretion of peptidases. **(A)** Fluorescence microscopy images showing the location of diatom and bacterial cells under specified conditions and times. Images include Brightfield, red chlorophyll autofluorescence, green SYBR Green-stained DNA, and a merged view of all channels. **(B)** Scatter plot showing diatom cell counts measured by flow cytometry, with lines representing the average of three replicates. Colors distinguish different media treatments. **(C)** Box plot showing the halo diameter generated on casein plates by different filtrates. A t-test was used to compare treatments, with significance indicated by stars.

To determine whether the presence of *T. pseudonana* was required to trigger the secretion of algicidal compounds, we attempted to grow *T. pseudonana* in exudates collected from (i) diatom mono-cultures, (ii) *A. macleodii* EP mono-cultures grown in diatom media, and (iii) co-cultures, all supplemented with MB. As expected, the growth rate of *T. pseudonana* in its mono-culture exudate was not significantly different from growth in mono-culture media (**(Fig. 4B).** In contrast, *T. pseudonana* did not grow in either the *A. macleodii* EP mono-culture exudate supplemented with MB or the co-culture exudate supplemented with MB (ANOVA followed by post hoc pairwise comparisons with Bonferroni-adjusted p-values: Control - Diatom Exudate=1, Control - Bacteria Exudate <.0001, Control - co-culture+MB Exudate <.0001, Bacteria Exudate - co-culture+MB Exudate=1) **(Fig. 4B)**. These results implied that (1) the presence of diatoms is not required to induce the algicidal response in *A. macleodii* EP, and (2) the algicidal agent is secreted into the media by the bacteria.

Motivated by the observation that peptidases changed their expression over the course of the experiment **(Fig. 3D-Fig. S6A)** and that the PSORTb algorithm [44] predicted that approximately 50% of the peptidases are extracellular, we used casein plates to test whether active peptidases or proteases were secreted into cell-free exudates. The presence of an active peptidase or protease within the exudate will cause the white color of the plate to fade, generating a measurable halo. Cell-free exudates from the *A. macleodii EP* mono-culture or co-cultures supplemented with MB both prevented diatom growth and generated casein halos indicating the presence of active proteases or peptidates **(Fig. 4C-Fig. S5A)**. The cell-free exudates from exponential co-cultures without the MB supplement did not generate a halo, despite the bacteria having the highest transcript levels associated with peptidases. This was most likely due to the low concentrations of *A. macleodii* EP in these co-cultures, presumably due limited availability of organic matter and the diatom defensive response **(Fig. 3D-4C-Fig. S5A)**.

### A. macleodii growth-motility trade-off is a conserved regulatory feature, but its directional progression in co-culture is host-specific

To determine whether *A. macleodii* EP displayed similar transcriptional patterns regardless of the co-culture host, we reanalyzed publicly available RNA-seq data from a co-culture experiment of *A. macleodii* MIT1002 with the marine photosynthetic bacterium *Prochlorococcus* NATL2A [45]. The new set of reads was aligned to the Te101 genome [35] to allow a comparison of genes between the two experiments.

The *A. macleodii* EP transcript levels profile in co-culture with *T. pseudonana* displayed an opposite pattern to that of *A. macleodii* MIT1002 in co-co-culture with *Prochlorococcus* NATL2A: those genes transcribed at high levels at the 30min time point when co-cultured with *T. pseudonana* were transcribed at high levels at the 12, 24, and 48 Hrs time points when co-cultured with *Prochlorococcus*, and vice versa **(Fig. S6)**. To further characterize this pattern, we focused those genes encoding ribosomal proteins and chemotaxis-related proteins **(Fig. 3E-G)**. The fraction of transcripts encoding ribosomal proteins increased over time in co-culture with *T. pseudonana,* whereas this fraction decreased over time in co-culture with *Prochlorococcus* **(Fig. 5A)**. Genes encoding chemotaxis proteins displayed the opposite pattern: the fraction of transcripts encoding chemotaxis proteins decreased over time in co-culture with *T. pseudonana,* whereas this fraction increased over time in co-culture with *Prochlorococcus* **(Fig. 5B)**. Finally, we calculated the ratio between the fraction transcripts encoding ribosomal proteins and chemotaxis-related proteins as a proxy for either cell growth or motility. Based on this proxy, *A. macleodii* EP shifts from a motility to a growth profile in co-culture with *T. pseudonana*, whereas *A. macleodii* MIT1002 shifts from a growth to a motility profile in co-culture with *Prochlorococcus* **(Fig. 5C)**. We conclude that the progression of gene expression in co-cultures is host-dependent and potentially strain-dependent, and that *A. macleodii* adjusts the growth-to-motility ratio in response to its environment.

**Figure 5.**
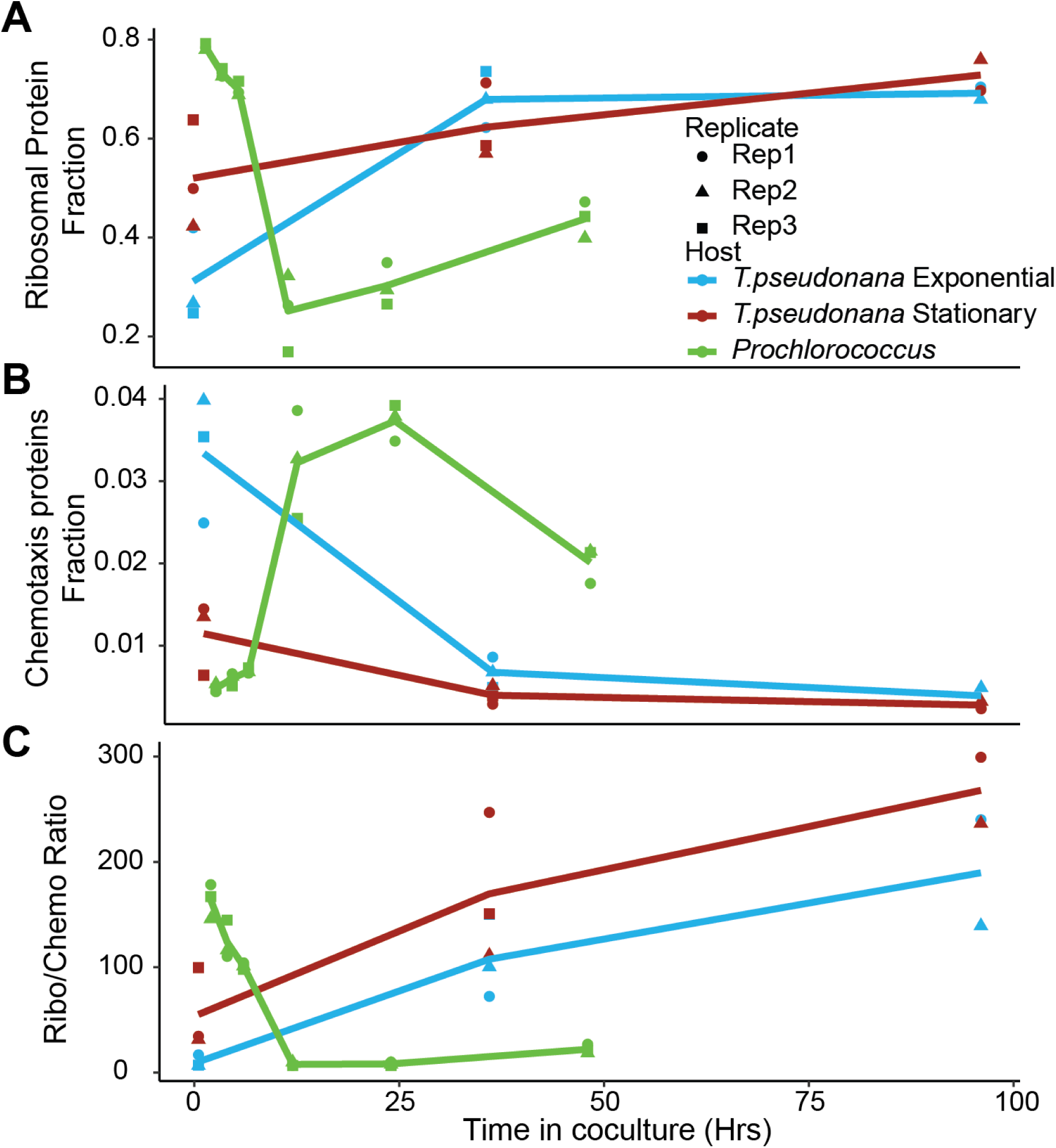
*A. macleodii* expression progression in co-culture is host-dependent, fluctuating between motility and growth. Scatter plots showing ribosomal proteins **(A)**, chemotaxis-related genes **(B)**, and the ratio between them **(C)**, representing their fraction of transcription in co-culture with T. pseudonana (Exponential—blue, Stationary—magenta) and Prochlorococcus (green). Replicates are indicated by different symbol types (circles, triangles, squares), with lines representing the average of two or three replicates.

## Discussion

In this study, we explored how diatom growth phase physiology influences interactions with bacteria. We discovered a dynamic interaction between a recently isolated Equatorial Pacific strain of *A. macleodii* and the model diatom *T. pseudonana*. Monitoring co-cultures, we observed bacterial growth only when *T. pseudonana* was in the stationary phase or when an external substrate source was provided. When bacteria grew to at least 1 × 10⁶ cells/mL, they exhibited an algicidal effect on *T. pseudonana*. We propose a two-stage model of pathogenic progression: initially, *A. macleodii* EP releases peptidases and/or proteases that compromise diatom cell structures, leading to cell lysis; subsequently, bacterial clusters accumulate around the resulting diatom debris. Moreover, we show that *A. macleodii* EP growth is limited in the presence of diatom cells, suggesting that the diatom generates an active defense response **(Fig. 6-upper)**. Similar to previous studies [7], our model highlights how the presence of sufficient substrate triggers the algicidal effect of *A. macleodii* EP. A key difference between the two strains is that, in the previous study, *A. macleodii* transitioned from mutualism to weak parasitism during co-culture with exponentially growing *T. pseudonana* cells, with the algicidal effect only triggered by the addition of an external substrate. In contrast, the EP strain was unable to grow when co-cultured with *T. pseudonana* in exponential phase due to the diatom’s defensive response. However, when the co-culture was initiated with diatom cells in stationary phase or with an external substrate source, the EP strain overcame the growth limitation and exhibited a significant algicidal effect. Moreover, both strains show similarities and differences in their algicidal mechanisms. Although both likely rely on peptidases/proteases for the algicidal effect, the EP strain does not require attachment. Co-cultures of additional *A. macleodii* strains with algae have shown competition for nitrate when co-cultured with the pennate diatom *Phaeodactylum tricornutum* [16], and a mutualistic interaction with the haptophyte *Isochrysis galbana* [14]. Together, these results underscore the complexity and context-dependence of bacteria-diatom interactions, highlighting how host physiology and environmental factors jointly shape microbial relationships in marine ecosystems.

**Figure 6.**
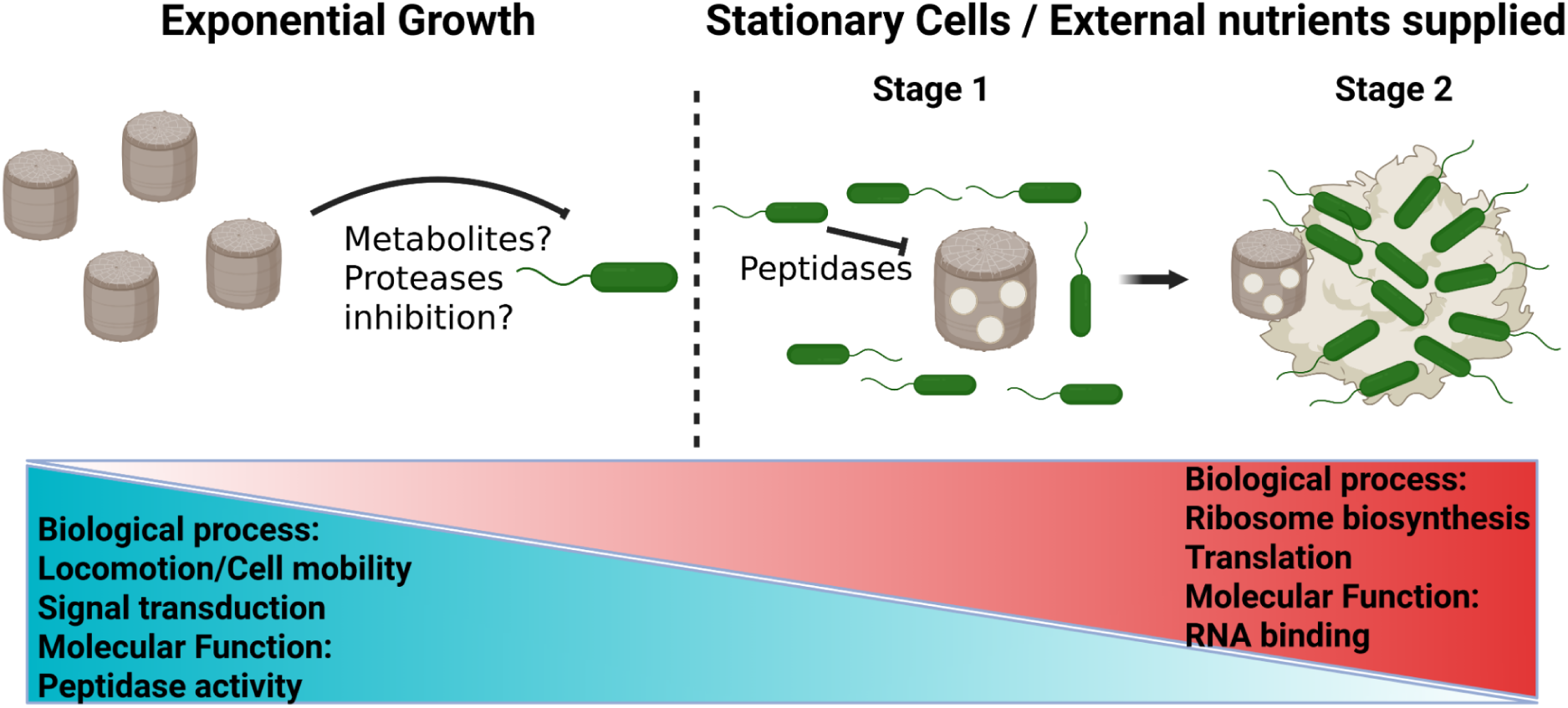
*A. macleodii* EP and *T. pseudonana* interaction regimes model. The initial conditions of *T. pseudonana* culture determine the outcome of its interaction with *A. macleodii* EP. During exponential growth, the diatom appears to control bacterial proliferation; however, in the stationary phase, or if external nutrients are supplied, bacterial cells thrive and exhibit an algicidal effect. This effect unfolds in two stages: first, bacterial growth accompanied by the secretion of a cocktail of peptidases, and subsequently, the formation of cell clusters around diatom debris. In response, *A. macleodii* EP senses its environment and adjusts the expression of genes involved in key biological functions to suit the prevailing conditions.

The transcriptomic landscape of *A. macleodii* EP across *T. pseudonana* growth phases reveals how the bacteria adapts to the diatom’s physiological state. In response to different diatom stages, *A. macleodii* EP modulates key gene groups, transitioning from a motility and sensing profile—characterized by locomotion, signal transduction, and peptidase activity—to a growth-oriented profile involving ribosome biosynthesis, translation, and RNA binding **(Fig. 6-lower)**. These shifts suggest that *A. macleodii* EP balances motility and growth depending on diatom state, likely reflecting stage-specific ecological needs. This pattern is consistent with a previously proposed trade-off between bacterial motility and growth, wherein the energetic and proteomic costs of maintaining motility machinery constrain investment in biosynthetic processes [46, 47].. This energetic and proteomic cost can divert resources away from growth, especially in environments where motility offers little benefit, such as well-mixed or nutrient-rich conditions [46, 47]. Previous studies in *E. coli* and other organisms have shown that translation is a limiting factor for growth [48–50], and the fraction of proteins allocated to the ribosome reflects the growth capacity. This likely explains why ribosomal protein expression fluctuates over time and across diatom physiological states in response to changes in substrate availability. In contrast, we observed that the chemotactic signaling machinery was upregulated at the beginning of the co-culture and in bacteria interacting with exponentially growing diatoms compared to those facing stationary diatoms **(Fig. 3G)**. Prior work has suggested that the *A. macleodii* algicidal effect and other bacteria’s attachment capacity to diatoms are dependent on chemotaxis [7, 51]. Additionally, it has been demonstrated that chemotaxis increases metabolic exchanges between heterotrophic bacteria and picoplankton [52]. In this study, we show that the algicidal effect of *A. macleodii* EP does not require cell-to-cell contact, as the extract alone is sufficient **(Fig. 4B)**. *A. macleodii* EP likely regulates its chemotaxis machinery depending on the availability of the substrates. The most concentrated areas of nutrient availability are located in the diffusive boundary layer or phycosphere surrounding phytoplankton cells, and *A. macleodii* EP could be chemotaxing towards an area of higher nutrient availability to support its growth when co-cultured with exponential-phase diatoms. Interestingly, independent of the downregulation of the chemotaxis machinery, bacterial cells facing exponential diatoms do not downregulate flagellar proteins, whereas cells facing stationary diatoms do **(Fig. 3G)**. The molecular phenotype is reflected in bacterial behavior, as cells facing stationary diatoms stop swimming and form aggregates around diatom debris **(Fig. 4A)**. This set of results demonstrates the long-lasting impact of the diatom’s growth phase at the beginning of co-culture on the bacterial response.

*A. macleodii* species is flexible in how it interacts with other microorganisms, and its interactions can range from pathogenic to mutualistic. Several strains of *A. macleodii* have been studied, and differences in their ecology have been inferred from their genomes and substrate utilization patterns [53, 54]. Our results and previous studies highlight that beyond genetic diversity, the outcome of diatom-bacteria interactions depends on the context in which both species interact, including substrate availability [7]. Bacterial transcriptomes respond to specific algal hosts [55] and nutrient conditions [56], enabling adaptation to changing environments. By comparing the co-culture gene expression profiles from this study to previous experiments with *Prochlorococcus* [45], we hypothesize that the host determines the *A. macleodii* gene expression progression in co-culture **(Fig. 5 - Fig. S6)**. A major difference between the two experiments is that while *A. macleodii* EP shifts from a motile to a growth expression profile in co-culture with *T. pseudonana*, a related strain follows the opposite trajectory in co-culture with *Prochlorococcus*. We cannot rule out the possibility that these differences are due to genetic content variations between strains (*EP* vs. MIT1002). Alternatively, the observed differences may result from the fundamentally different nature of each interaction. In co-culture with *Prochlorococcus*, *A. macleodii* MIT1002 engages in a mutualistic relationship [45], whereas in co-culture with *T. pseudonana*, *A. macleodii* EP displays a pathogenic behavior. Moreover, the presence of *A. macleodii* MIT1002 facilitates *Prochlorococcus* survival in the dark by exchanging essential compounds, showing how the two species support each other’s metabolism [57, 58]. Additional studies have highlighted the collaborative role of *A. macleodii* in the ocean, showing that it shares siderophores to mobilize iron and extracellular enzymes to break down complex polysaccharides for community use [59, 60]. These findings suggest that in mutualistic contexts, such as with *Prochlorococcus*, *A. macleodii* is likely to receive growth-promoting cues from the host, triggering a switch to a motile state only after the host enters stationary phase. In contrast, pathogenic interactions—such as those with *T. pseudonana*—initially induce a motile transcriptional profile, which may subsequently switch in response to signals triggered by the pathogenic effect. Interestingly, even when co-cultured with substrate rich stationary-phase diatoms, *A. macleodii* EP still shifts from motility to growth transcriptomic profiles **(Fig. 5)**, suggesting that the motile-to-growth transition may be part of an intrinsic host dependent response. We propose that modulating the balance between motility and growth is a key regulatory feature of *A. macleodii* interactions.

We discovered that the interactions of *A. macleodii* with phytoplankton vary depending on the growth phase of the host—a finding with wide-ranging ecological implications, such as potential roles in regulating algal blooms, reshaping microbial community composition, and enhancing carbon export through aggregate formation. Phytoplankton blooms exhibit a similar growth pattern as cultures, with an exponential growth phase followed by a demise after reaching maximum cell abundance [61]. It has been well established that bacterial composition changes throughout the algal blooms [24, 62]. Our results suggest that algae may limit the growth of pathogenic bacteria during the expansion phase of a bloom, but not once maximal algal growth has been reached—similar to what we observed in our co-culture experiments **(Fig. 1A)**. Additionally, examples of bacterial-mediated protection of unicellular algae against algicidal bacteria have been documented [11]. The effect of interactions with other bacteria on the algicidal pattern of *A. macleodii* is unknown; however, in high nutrient conditions, copiotrophic *A. macleodii* might be able to outcompete other bacteria and reach high enough concentrations to be responsible for bloom demise. Additionally, particle aggregation increases as a consequence of algal blooms [63, 64]. The ability of Alteromonas species to form aggregates through the secretion of sticky exopolymers has been demonstrated [65], and at least one of the genes required for their production—*UDP-glucose 4-epimerase*—is conserved across *A. macleodii* strains [53]. We observed increased aggregate formation at the 8-day timepoint, which coincided with a significant accumulation of *UDP-glucose 4-epimerase* transcripts over time **(Fig. S7)**. Notably, transcript levels did not significantly differ between stationary and exponential co-cultures, suggesting that the differences in aggregation are likely driven by higher cell densities in coculture with diatoms at stationary phase, rather than differences in gene expression per se **(Fig. S7)**. The formation of aggregates by *A. macleodii* could affect microbial interactions and carbon cycling: larger aggregates are more likely to be grazed by larger zooplankton, facilitating nutrient transfer up the food web or their increased size can accelerate sinking rates, potentially enhancing carbon export from the surface to the deep ocean [55].

A key finding of our study is that the presence of *T. pseudonana* limits *A. macleodii* EP cell division. The molecular mechanism behind this effect remains to be discovered. One possibility is the secretion of metabolites that limit bacterial growth. It has been previously shown that diatoms secrete specific compounds to control bacterial companions [13, 66], and this is a plausible mechanism in this interaction, as it does not require direct cell-to-cell contact. Analyzing the secreted metabolome of co-cultures, compared to mono-cultures, will help identify potential regulatory metabolites. Alternatively, the diatom could secrete enzymes that limit bacterial growth. Among the limited number of differentially expressed genes, we detected three transcripts with predicted serine protease inhibitor activity—THAPSDRAFT_6004, THAPSDRAFT_36303a, and THAPSDRAFT_24702—which could potentially contribute to limiting bacterial pathogenicity and their ability to extract resources from the diatom by truncating the function of bacterial proteases [67] **(Supplemental Table 4, Fig-lower))**. Additionally, it remains unclear how cells regulate the activation of this mechanism. The differences in bacterial growth between mono-culture and co-culture exudates suggest that the defensive response is not constitutive **(Fig. 2C-Fig. S3C),** implying that the bacterial presence triggers a molecular cascade that culminates in the defensive effect. This type of communication has been described in other organisms, although it remains poorly understood in diatoms [68]. We detected a Natriuretic peptide receptor (THAPSDRAFT_263505), predicted to be located in the membrane, and a Peripheral-type benzodiazepine receptor (THAPSDRAFT_2880), predicted to be located in the mitochondria, both of which were differentially expressed and could potentially be associated with triggering the signaling cascade. Additionally, a Serine/threonine protein kinase (THAPSDRAFT_1388) and a Serine/threonine protein phosphatase (THAPSDRAFT_36303) were differentially expressed and may be involved in the propagation of the signal **(Supplemental Table 4)**. As a potential consequence of the signaling cascade, we observed a significant fraction of differentially expressed genes associated with chromatin structure and dynamics kogClass **(Supplemental Table 4; Fig. S8C)**. Ten histones were downregulated in coculture with the bacteria, suggesting a major chromatin reconfiguration **(Supplemental Table 4; Fig. S8C)**. A major caveat in this analysis is that ∼32% of the differentially expressed genes remain unannotated, highlighting how a substantial portion of diatom biology is still understudied **(Supplemental Table 4; Fig. S8C)**. Additionally, there is evidence in other organisms of post-transcriptional [69, 70] and post-translational [71, 72] responses to bacterial presence. Further studies on post-transcriptional regulation may provide insights into the mechanisms underlying *T. pseudonana*’s defensive response.

Co-culturing *A. macleodii* EP with *T. pseudonana* revealed that bacterial transcriptional programs and phenotypes are modulated by host growth phase, substrate availability, and inducible defense responses. These findings build on prior work linking phytoplankton physiology to microbial community structure, and extend it by identifying molecular mechanisms through which host state regulates bacterial behavior. How growth phase-driven regulatory dynamics influence ecosystem-scale processes such as bloom collapse, particle aggregation, and carbon export in marine environments remains a critical area for further study.

## Acknowledgements

We would like to thank the members of the Armbrust Lab for insightful discussions, particularly Megan Schatz for her training and assistance with growing algal cultures. D.W. is supported by postdoctoral fellowships from the European Molecular Biology Organization (EMBO-ALTF 62-2022) and the Human Frontier Science Program (HFSP #982011). R.D.’s summer research was funded by the Cooperative Institute for Climate, Ocean, and Ecosystem Studies (CICOES, 2024). This work was supported by grants from the Simons Foundation (#721244, EVA) and the National Science Foundation (NSF #2051212).

## Ethics declarations

### Competing interests

The authors declare no competing interest.

## Notes

### Competing Interest Statement

The authors have declared no competing interest.

