## Supplementary Figures for "Growth phase-specific gene regulation and algicidal interactions between a new *A. macleodii* strain and the model diatom *T. pseudonana*"

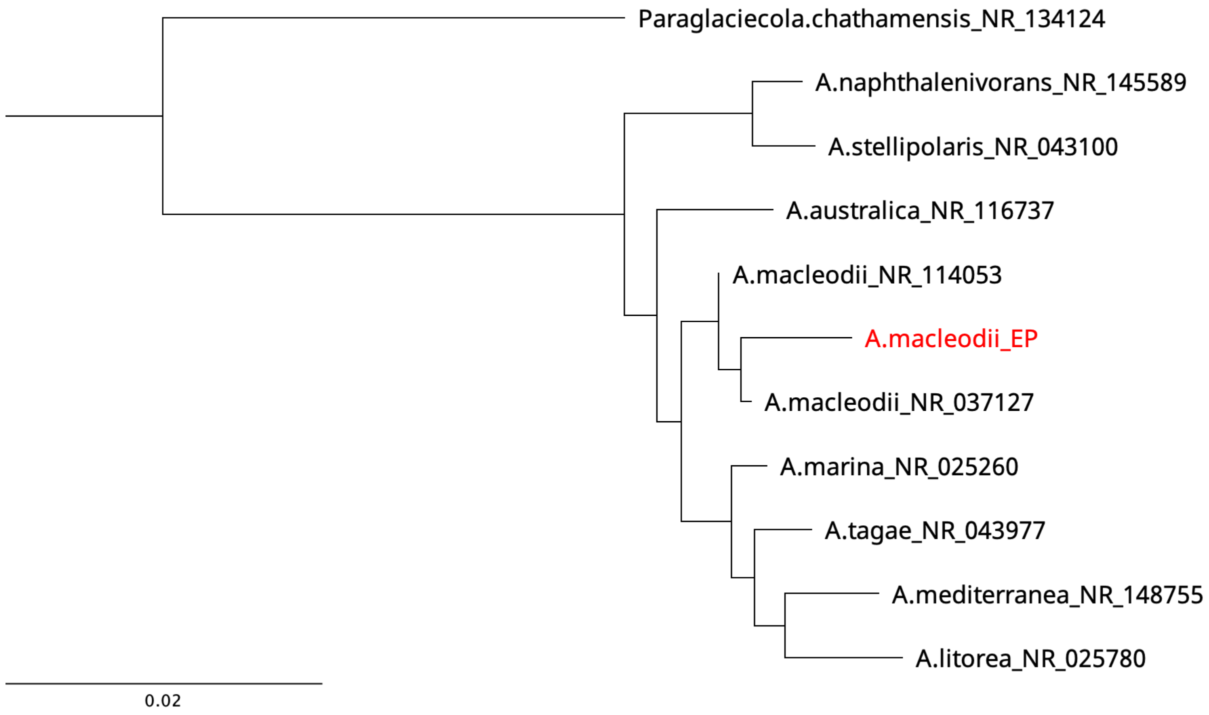


**Supp 1. New Equatorial Pacific bacterial strain cluster within *A. macleodii* species.** A maximum likelihood 16S rRNA phylogenetic tree of *A. macleodii* EP and closest relatives based on NCBI BLAST search. The tree is rooted in the outgroup *Paraglaciecola chathamensis*.

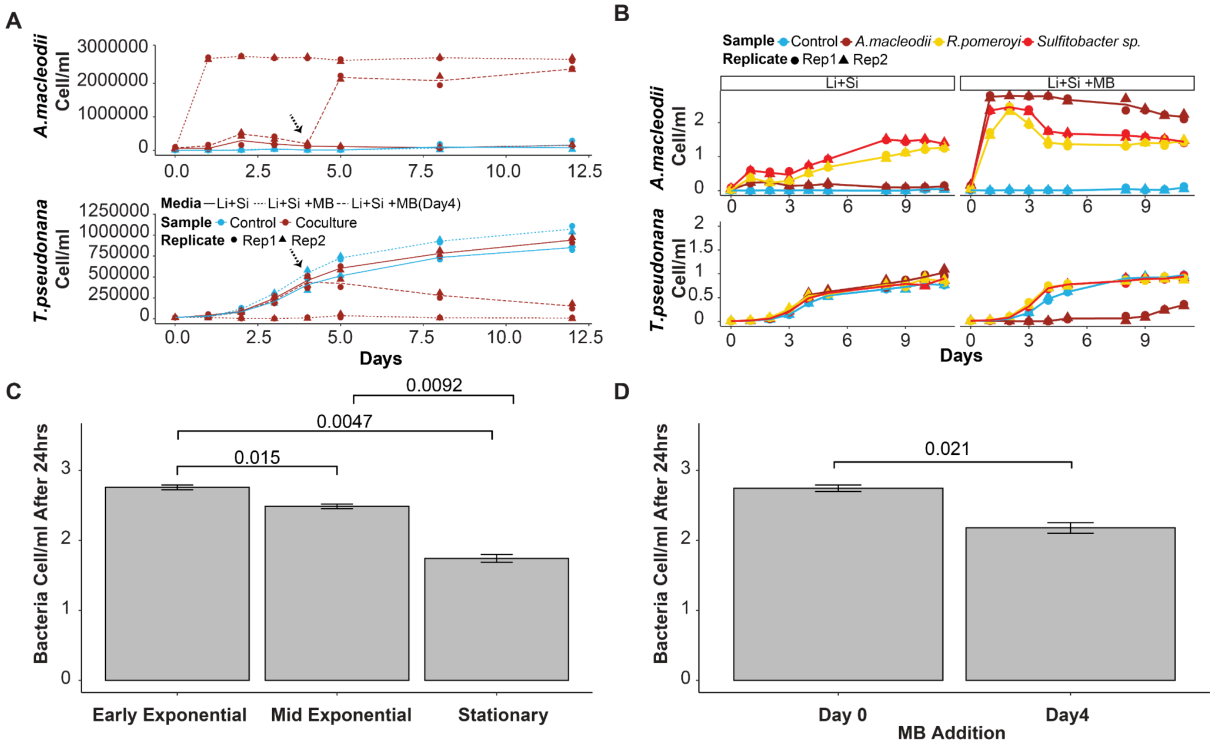


**Supp 2. *T. pseudonana* growth phase and substrate supply determine interaction dynamics with *A. macleodii* EP. (A)** Similar to **Fig. 1A**, showing cell counts in co-culture (magenta symbols) or mono-culture (blue symbols). Dashed lines indicate the timing of early and late MB addition. **(B)** Similar to **Fig. 1A.**. Each facet represents cultures grown in L1+Si with or without MB supplementation. Colors indicate specific cocultures: *A. macleodii* (magenta), *R. pomeroyi* (yellow), *Sulfitobacter* sp. (red), and control (blue symbols). **(C)** Bar plot showing bacterial cell counts (cells/mL) 24 hours after coculture initiation across different diatom growth phases. Wilcoxon signed-rank test p-values are indicated for each comparison. **(D)** As in (C), comparing early vs. late MB addition to the coculture.

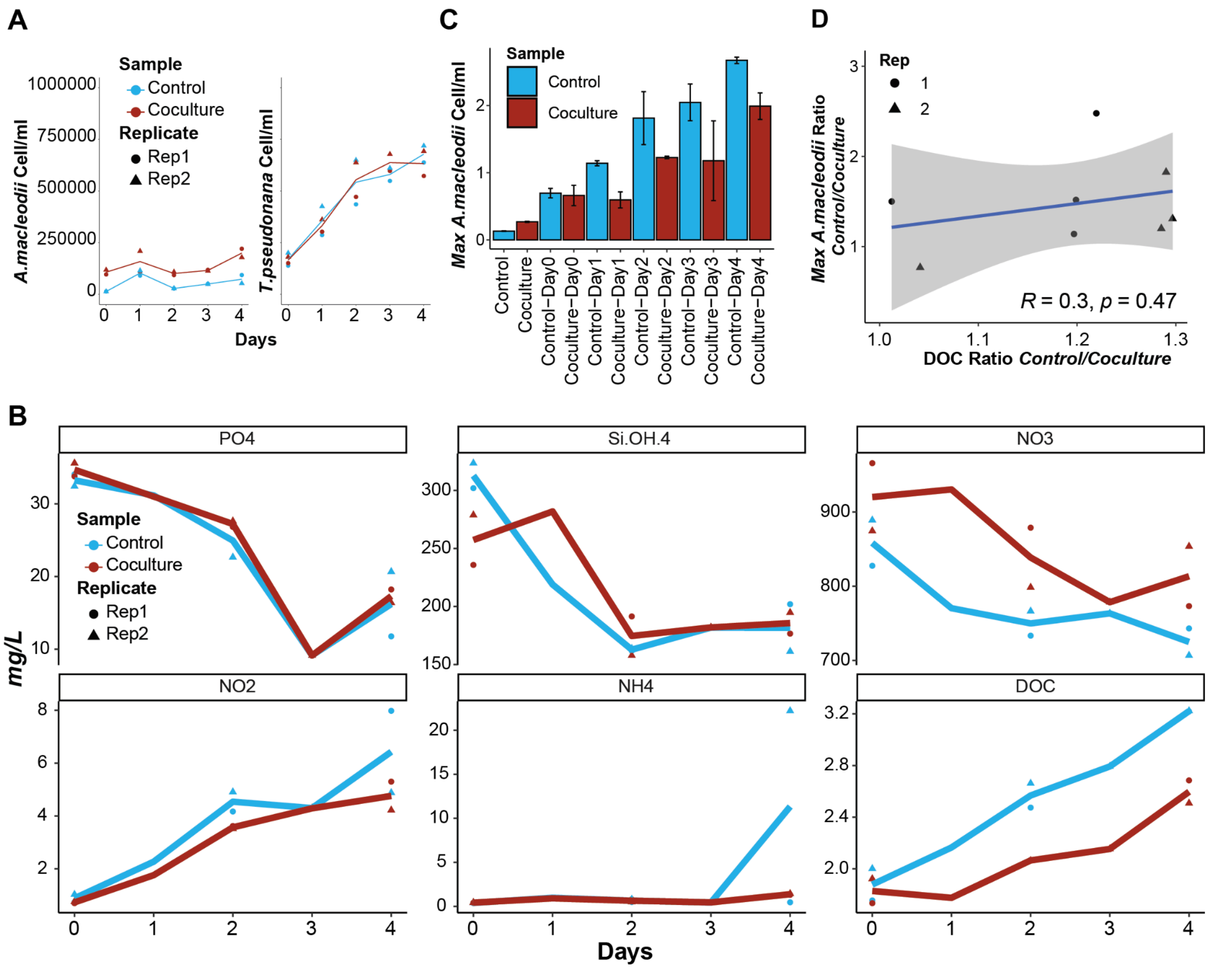


**Supp 3. *T. pseudonana* imposes constraints on bacterial growth beyond nutrient limitation. (A)** Similar to **Fig.1A**, at mid-exponential stage. **(B)** Scatter plots showing the selected nutrient concentrations of the corresponding day's exudates. Color code distinguishes control from coculture. **(C)** Bar plot showing maximal bacterial cell counts (cells/mL) in the original samples and in each exudate. The error bar shows the standard deviation, and the color code distinguishes control from coculture samples. **(D)** Scatter plot between the ratio of maximal bacterial cell counts between the Control and Coculture, plotted against the DOC ratio. The Pearson correlation coefficient and associated p-value are included.

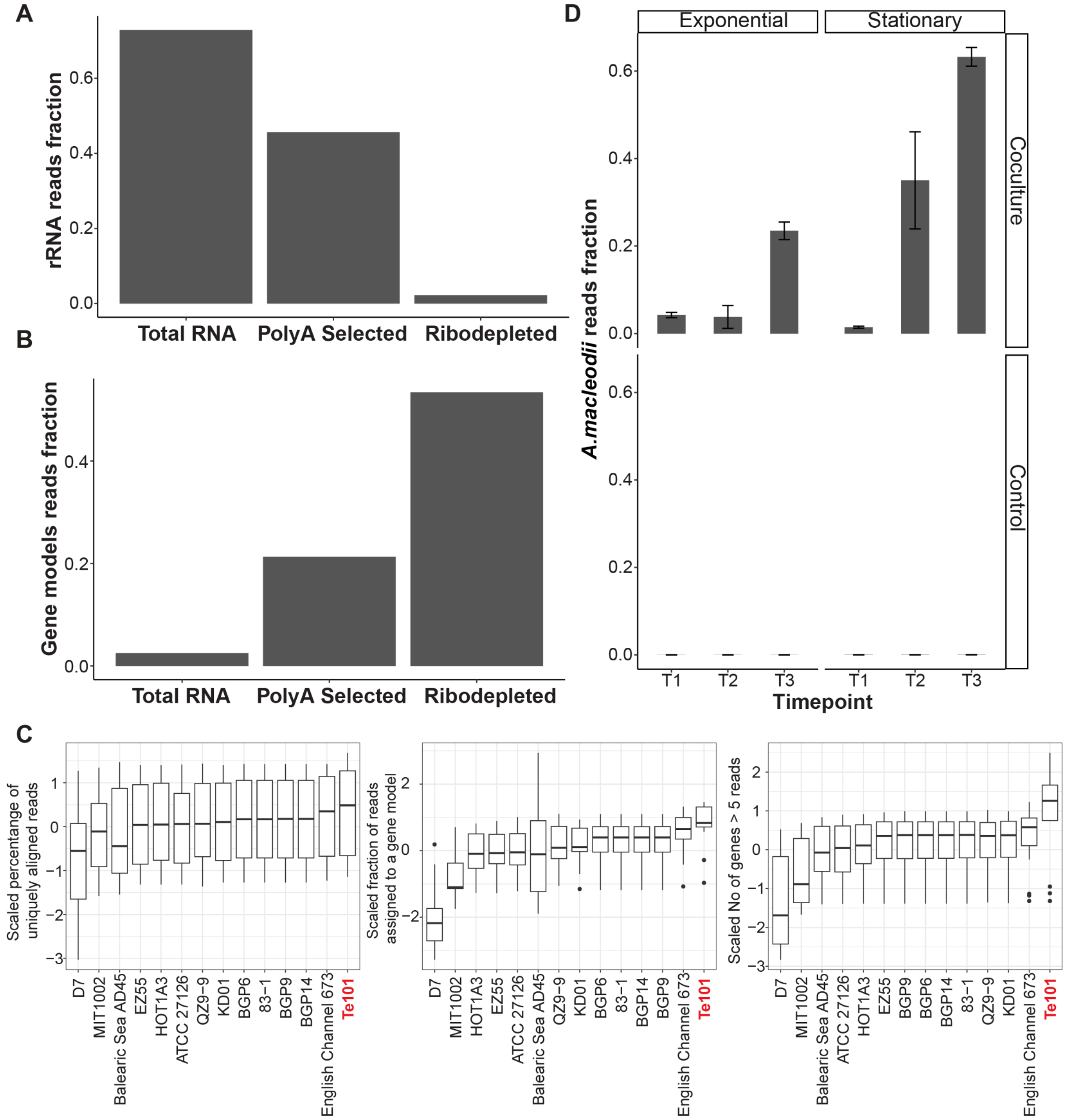


**Supp 4. Tailored Duo-RNAseq protocol efficiently removes *T. pseudonana* rRNA and accurately assigns reads to the corresponding genome. (A)** Barplot showing the rRNA reads fraction in three different *T. pseudonana* RNA library preparation protocols (Total RNA, Poly(A) selection, and Ribodepletion). **(B)** Similar to A, but for the gene models' read fraction. **(C)** Boxplot showing the percentage of uniquely aligned reads (left), the fraction of reads assigned to a gene model (middle), and the number of gene models with more than five reads for all coculture samples per *A. macleodii* genome (Table 2). Each metric was scaled across coculture time points. The selected genome is highlighted in red. **(D)** Barplot showing the fraction of A. macleodii EP-assigned reads at each time point. Error bars represent the standard deviation of three replicates. Facets distinguish between Control and Coculture, as well as Stationary and Exponential samples.

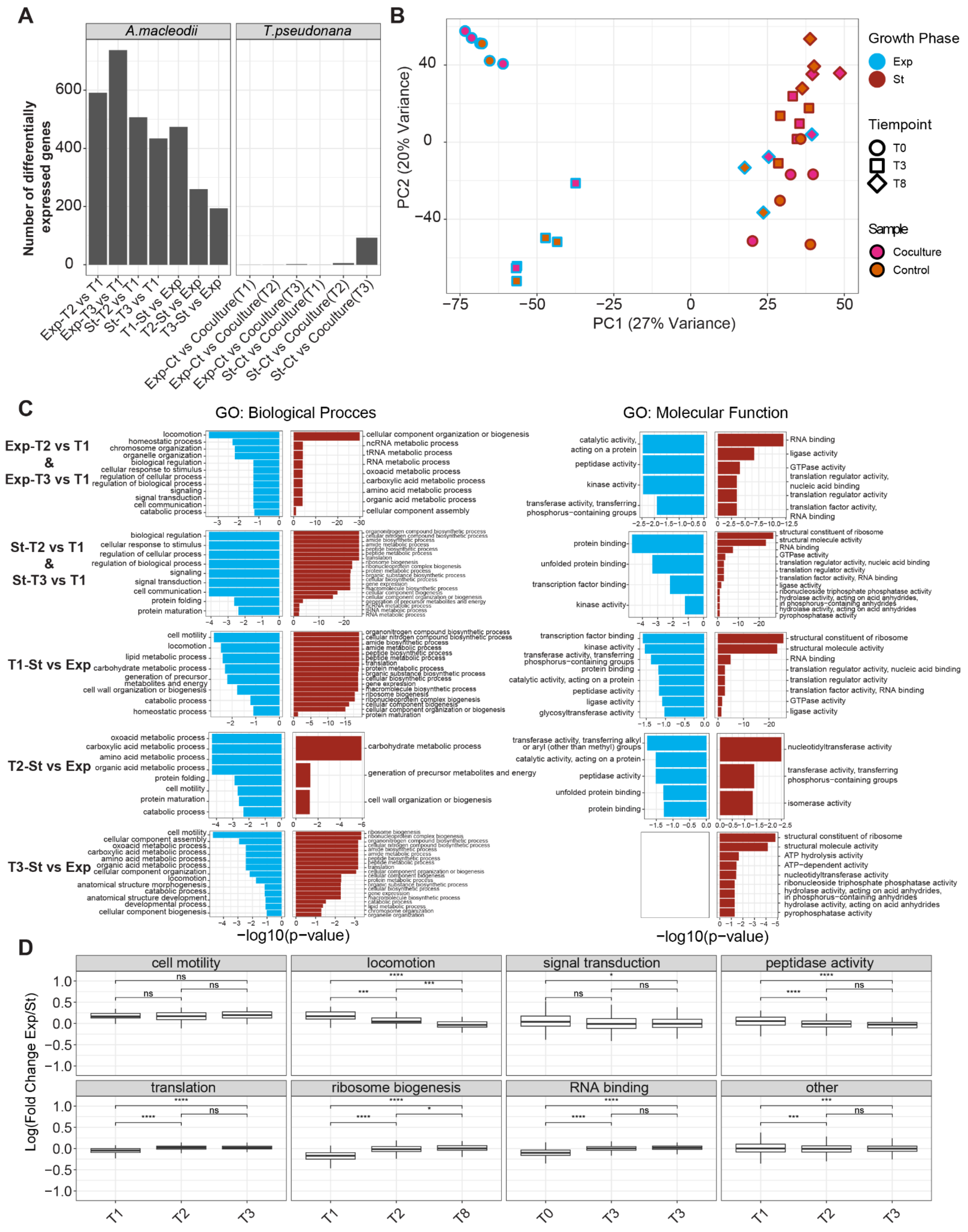


**Supp 5. Gene expression regulatory landscape in the *A. macleodii* EP–*T. pseudonana* interaction. (A)** Bar plot showing the number of differentially expressed genes in both genomes for specific comparisons**. (B)** Principal component analysis (PCA) using scaled RPKM reads from *T. pseudonana*. The internal color code corresponds to the control (orange) and coculture (pink), while the external color code represents the initial diatom growth phase (exponential - blue, stationary - magenta). Shapes represent the time points: Circles for T1, Squares for T2, and Triangles for T3. **(C)** Gene ontology terms associated with upregulated (magenta) or downregulated (blue) genes in the indicated comparisons. **(D)** Log fold change between Exponential and Stationary for the selected GO at each time point. Significance of the Wilcoxon signed-rank test p-values are indicated for each comparison.

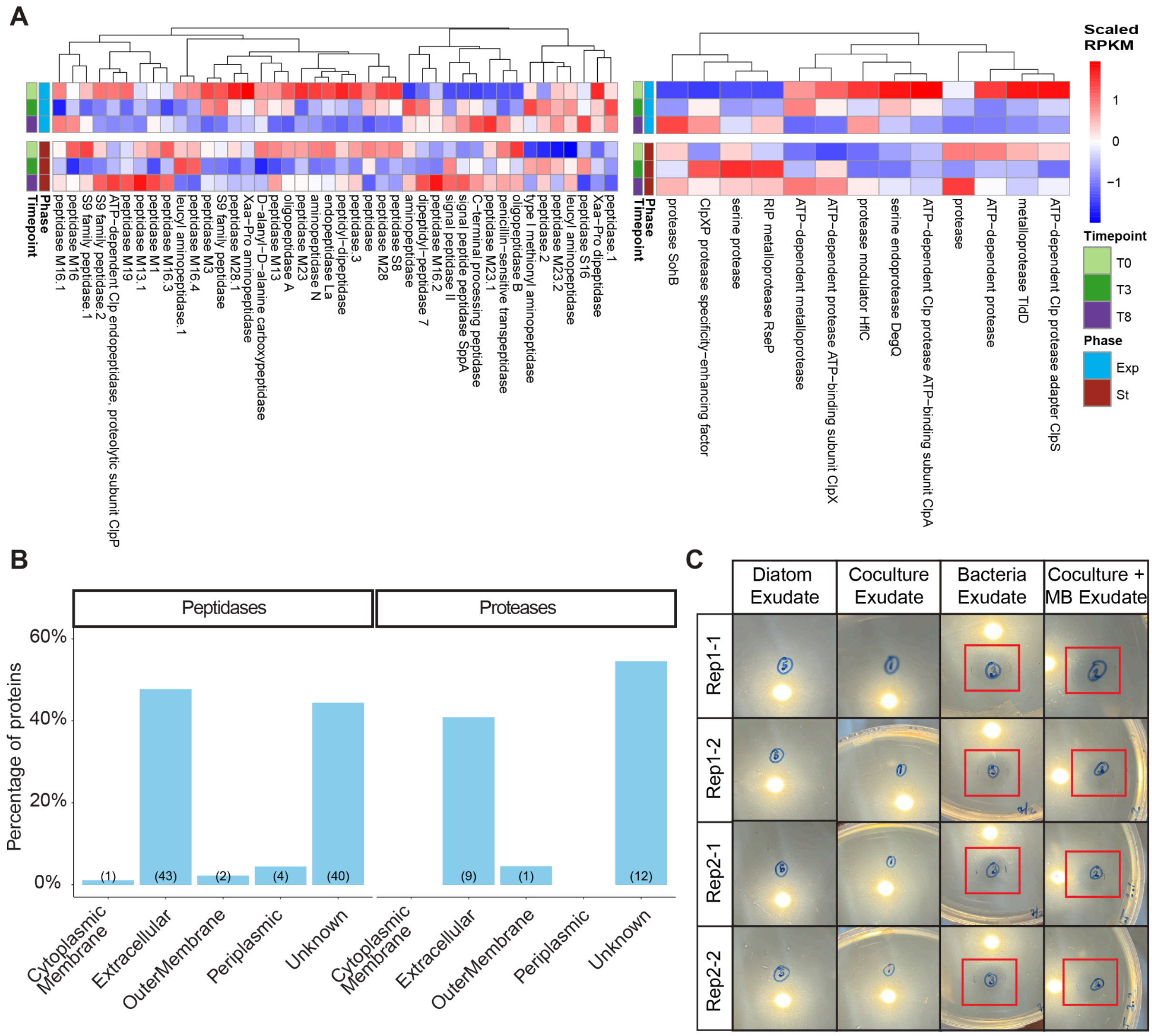


**Supp. 6. *A. macleodii EP* secretes peptidases or/and proteases during the coculture. (A)** Heatmaps of gene expression for *A. macleodii* peptidases (left) and proteases (right). The color scale represents scaled TMM-normalized RPKM values. Columns (genes) were clustered using Euclidean distance with the pheatmap function in R. Row annotations indicate sample type: exponential phase (blue) and stationary phase (magenta). Time points are marked in light green (T1), green (T2), and purple (T3). For genes with identical annotations, a dot and a number were appended to the gene name. **(B)** Bar plot showing PSORTb localization prediction of peptidases (left) and proteases (right) encoded in the *A. macleodii* Te101 genome. The y-axis represents the percentage of proteins in each localization category, and the numbers in parentheses indicate the absolute number of proteins per localization. **(C)** Table showing photographs of the casein agar plates after 6 days had elapsed. The left legend indicates the replicant group, while the top legend indicates the condition. Halos (clear circles) indicate the presence of proteases; the halos are highlighted with red boxes.

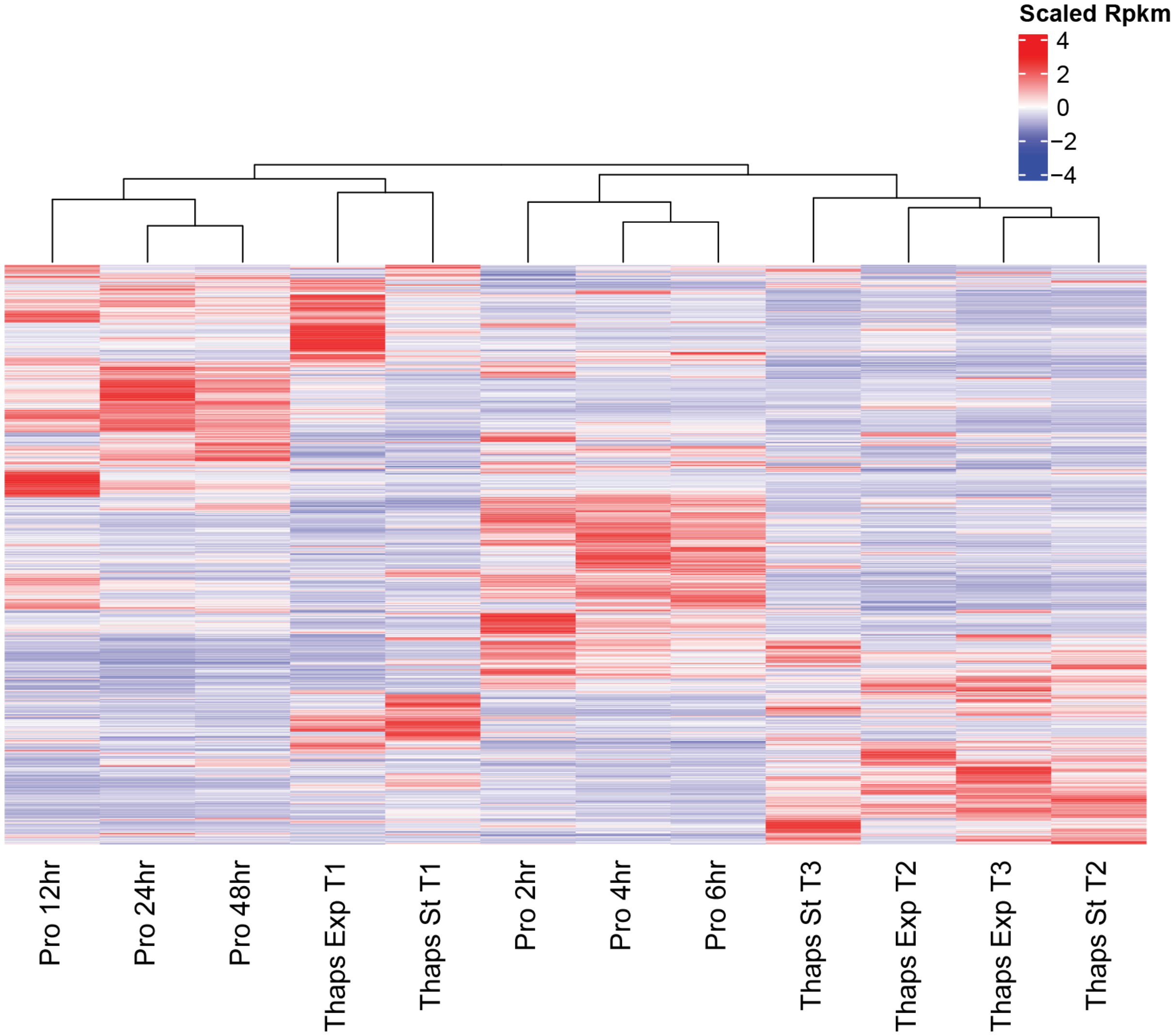


**Supp. 7. *A. macleodii* Expression profiles cluster according to time in coculture and host.** Heatmap of gene expression across both co-cultures. The color scale represents scaled TMM-normalized RPKM values. Rows and columns were clustered using Euclidean distance with the pheatmap function in R.

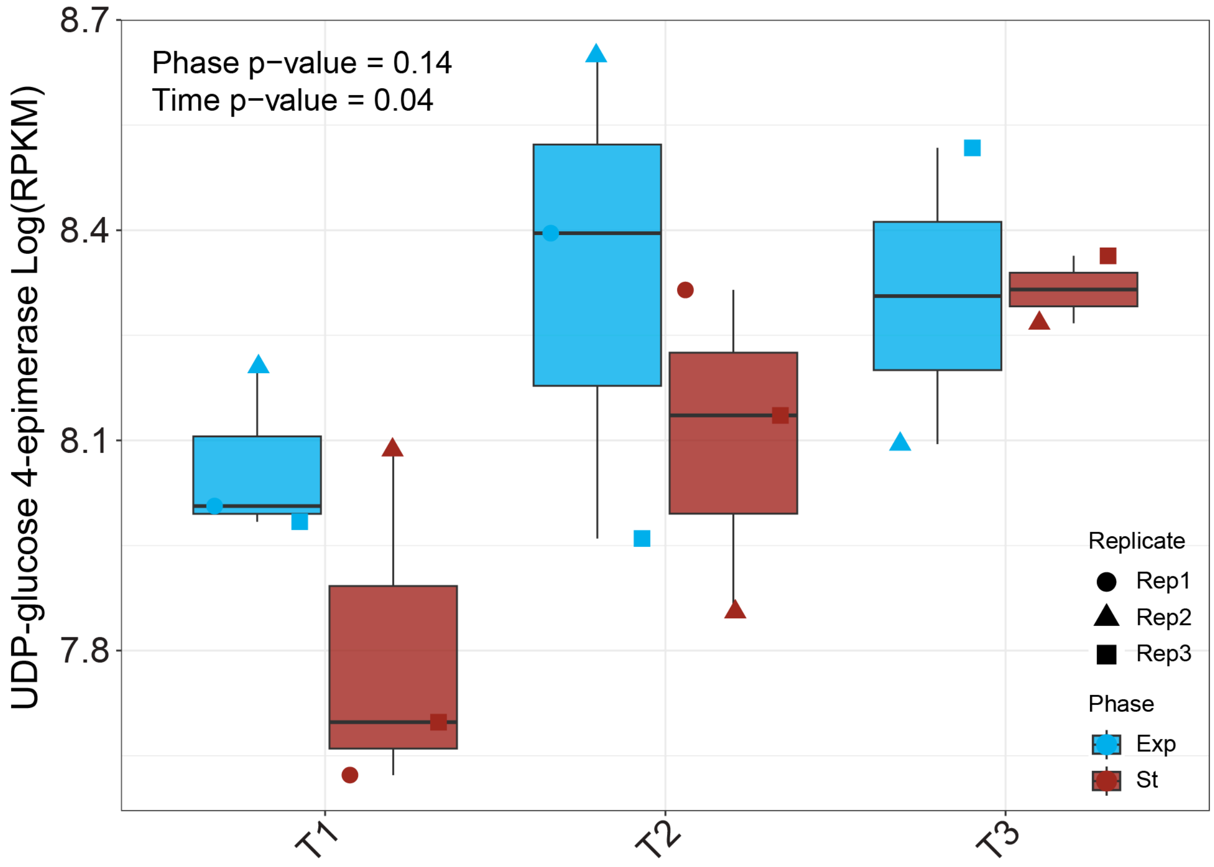


**Supp. 7. *UDP-glucose 4-epimerase* transcript levels increase with time in coculture.** Boxplots showing the RPKM values of the *A. macleodii* gene *UDP-glucose 4-epimerase* in exponential (blue) and stationary (magenta) phases of diatom cocultures. Each biological replicate is represented by a dot with a distinct shape.

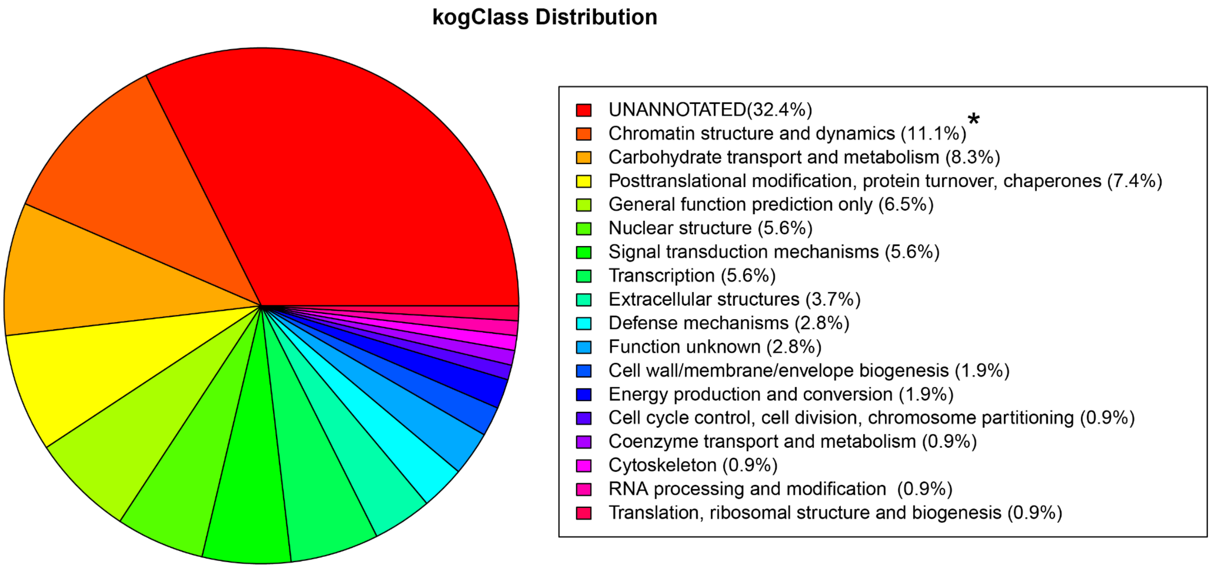


**Supp. 8. Chromatin structure and dynamics genes are overrepresented among *T. pseudonana* differentially expressed genes.** Pie chart showing the distribution of KOG class categories among differentially expressed genes. * indicates statistically significant enrichment compared to the full transcriptome (Chi-squared test for proportions).
