## Supplemental Tables for "Growth phase-specific gene regulation and algicidal interactions between a new *A. macleodii* strain and the model diatom *T. pseudonana*"

**Supplemental Table 1:** Species used in this study.

**Supplemental Table 2.** Blast results of comparing *A. macleodii* 16S to NCBI references. Top 30 references based on e-value and similarity score.

**Supplemental Table 3:** *A. macleodii* genomes list.

**Supplemental Table 4:** List of differentially expressed genes in *T. pseudonana.*

**Supplemental Data 1:** Merged forward and reverse Sanger 16S sequences for *A. macleodii* EP.
