## Supplemental Data 1 for "Growth phase-specific gene regulation and algicidal interactions between a new *A. macleodii* strain and the model diatom *T. pseudonana*"

>A.macleodii_EP_16S

NNNNNNNNNNNNNNNNNNCNNNNNNNNNNNTGNNNNNNNNNNNNNNNNNNNNNNTNNNNNNNNNNNNNNTNNNNNNNNNNNNNN
NNNNAGTGTNNNNNNNNNNNNNNNNNNNGNNNCNGNNNNCNNNNGNCNNNNNACTNNNTNNNNANNNNNTNNNNNTNNNN
NCNANGNCNNTANANNTNNNNNNNNNTNNNNGNGGTNAAANNNNNNCCCCCCCCCNTNNNNNNNTTTTTNNNNCTGCTCAG
ATTGACGCTGGCGGCAGGCCTAACACATGCAAGTCGAACGGTAACATTTCTAGCTTGCTAGAAGATGACGAGTGGCGGACG
GGTGAGTAATGCTTGGGAACTTGCCTTTGCGAGGGGGATAACAGTTGGAAACGACTGCTAATACCGCATAATGTCTTCGGAC
CAAACGGGGCTTCGGCTCCGGCGCAAAGAGAGGCCCAAGTGAGATTAGCTAGTTGGTAAGGTAACGGCTTACCAAGGCGA
CGATCTCTAGCTGTTCTGAGAGGAAGATCAGCCACACTGGGACTGAGACACGGCCCAGACTCCTACGGGAGGCAGCAGTG
GGGAATATTGCACAATGGGGGAAACCCTGATGCAGCCATGCCGCGTGTGTGAAGAAGGCCTTCGGGTTGTAAAGCACTTTC
AGTTGTGAGGAAAAGTTAGTAGTTAATACCTGCTAGCCGTGACGTTAACAACAGAAGAAGCACCGGCTAACTCCGTGCCAG
CAGCCGCGGTAATACGGAGGGTGCGAGCGTTAATCGGAATTACTGGGCGTAAAGCGCACGCAGGCGGTTTGTTAAGCTAG
ATGTGAAAGCCCCGGGCTCAACCTGGGATGGTCATTTAGAACTGGCAGACTAGAGTCTTGGAGAGGGGAGTGGAATTCCA
GGTGTAGCGGTGAAATGCGTAGATATCTGGAGGAACATCAGTGGCGAAGGCGACTCCCTGGCCAAAGACTGACGCTCATGT
GCGAAAGTGTGGGTAGCGAACAGGATTAGATACCCTGGTAGTCCACACCGTAAACGCTGTCTACTAGCTGTGTGTGTCTTTA
AGACGTGCGTAGCGAAGCTAACGCGCTAAGTAGACCGCCTGGGGAGTACGNCCGCAAGGTTAAAACTCAAATGAATTGACG
GGGGCCCGCACAAGCGGTGGAGCATGTGGTTTAATTCGATGCAACGCGAAGAACCTTACCTACACTTGACATGCTGAGAAG
TTACTAGAGATAGTTTCGTGCCTTCGGGAACTCAGACACAGGTGCTGCATGGCTGTCGTCAGCTCGTGTCGTGAGATGTTG
GGTTAAGTCCCGCAACGAGCGCAACCCTTGTCCTTAGTTGCCAGCCTTAAGTTGGGCACTCTAAGGAGACTGCCGGTGACA
AACCGGANGAAGGTGGGGACGACGTCAAGTCATCATGGCCCTTACGTGTANGGCTACACACGTGCTACAATGGCATTTACA
GAGGGAAGCGAGACAGTGATGTGGAGCGGACCCCTTAAAGANTGTCGTNNTCCNGGATTGNAGTCTGCAACTCGACTCCA
TGAAGTCNGGAGTCGCTAGTAATCGNNAGGTCAGANACTGNNGGGAATNCGTTCCNGGNNNTNNNANCNNCGCCACNNN
GGANNANGCNNGTGNGNTTCGANNNNGTANNANNNCGNGGCGNNNANNNNGANTATANNACNTTGATGCNNNNNANANN
GCNNNCNNCTCNTTGACNNNNNGNGNNTGNNNNNNNANTNNNNNNNNNNNNNGNNTTCNGNANNNNNANNCNGNNNNNN
NNGNNTNNNNANCNACNNNNNNNNNNNNNNNNNNGNGNNNNNNNNNNNANNNNNNNNNNNNNNNNNNNNNNNNNNNNNNNNTNA
NNNNGNNNNNNNNNNGANNNNCNNNNNNNNNNNNNNGGNNNNNNNNNNNNNNNNNNNNNNNNNNNNNNNNNNNNNNNNNNCNNNGNNN
NNNGGNNNN
